# Deadwood-related microhabitats in old-growth forests in Poland

**DOI:** 10.64898/2026.08.12.744401

**Authors:** Fabian Przepióra, Michał Ciach

## Abstract

Deadwood is a fundamental component of forest ecosystem, supporting biodiversity and driving multiple ecological processes. However, structures that may develop on downed coarse woody debris (CWD) create additional microhabitats that are used by numerous organisms and contribute to small-scale habitat heterogeneity. To date, quality of CWD is commonly characterized by its volume, diameter, tree species or stage of decomposition, while fine-scale structures occurring there remain not surveyed and their role in ecosystem is rarely quantified. Here, we introduce the novel concept of microhabitats on CWD. i.e. Deadwood-related Microhabitats (DreMs), defined as distinct features occurring on CWD that may provide shelter, breeding or foraging sites for forest-dwelling organisms. Using an original catalogue comprising 14 groups and 30 types, we inventoried DreMs on 6,003 CWD across 423 study plots located in best-preserved old-growth forests in Poland. We quantified the frequency and richness of DreMs and assessed the link between CWD characteristics and DreM richness in spruce, beech, willow-poplar, fir-beech and oak-lime-hornbeam forests. All inventoried DreMs occurred on both deciduous and coniferous taxa. The most frequent DreM included bryophyte mats, loose bark patches, insect galleries, fungal fruiting bodies and polypores. DreM richness increased with CWD diameter, more complex architecture and the presence of multiple decay classes within single debris. DreM richness peaked at intermediate decay classes and was higher on deciduous than on coniferous taxa. Our study is the first large-scale qualitative and quantitative assessment of DreMs in temperate forests. By focusing on forests characterized by ecological continuity and minimal human-related disturbance, the results provide a reference for downed deadwood-associated structures. By complementing inventory of tree-related microhabitats, DreMs extend potential monitoring schemes of habitat quality and contribute to biodiversity-oriented forest management.

**Highlights:** • Downed coarse woody debris (CWD) hosts Deadwood-related microhabitats (DreMs)

• Higher DreM richness is associated with CWD diameter

• DreM richness peak at intermediate stages of wood decay

• CWD of deciduous taxa support more DreMs than coniferous

• Diversified CWD and DreMs increase habitat heterogeneity for forest-dwelling taxa

## Introduction

Deadwood, including dead standing trees, fallen logs and uprooted trees, is recognized as a fundamental component of ecosystem (Gutowski et al., 2022). In temperate forests, deadwood supports a substantial proportion of forest-related biodiversity by providing substrate for saproxylic fungi, bryophytes, lichens, invertebrates and vertebrates (Stokland et al., 2012). The spatial distribution, diversity and long-term persistence of deadwood are, therefore, considered essential attributes of old-growth forests, influencing ecosystem resilience, nutrient dynamics and species persistence across multiple trophic levels (Harmon et al., 1986). Consequently, deadwood quantity and quality have become one of the key ecological indicators used in conservation, restoration and biodiversity-oriented forest management (Müller & Bütler, 2010, Lassauce et al., 2011).

Beyond its role as a substrate for specialized organisms, deadwood contributes to a wide range of ecological processes operating at different spatial scales. Large accumulations of deadwood regulate carbon and nutrient cycling (Harmon et al., 1986), influence water retention and hydrological dynamics (Błońska et al., 2018; Klamerus-Iwan et al., 2020), and shape microtopography through mound-and-pit systems associated with uprooted trees (Schaetzl et al., 1989). In riparian forests and stream ecosystems, fallen logs modify water flow and create important aquatic habitats (Gurnell et al., 1995). At broader spatial scales, high volumes of deadwood may also affect vertebrate behaviour and movement patterns by increased habitat heterogeneity and reduced visibility, thereby contributing to the so-called landscape of fear (Laundré et al., 2010). Such effects may alter habitat use by large herbivores and predators, ultimately influencing trophic interactions and forest regeneration (van Ginkel et al., 2019).

Downed coarse woody debris (CWD), i.e. fallen or uprooted trees and their large pieces – logs and branches, frequently functions as small-scale corridors facilitating dispersal of saproxylic organisms dependent on spatial continuity of woody substrates (Schiegg, 2000). Simultaneously, numerous vertebrates actively exploit deadwood-associated structures for shelter, movement, reproduction, thermoregulation or foraging (Carey & Johnson, 1995; Fritts et al., 2015; Phillips & Garrick, 2026). Crevices beneath loose bark or within decomposing wood are commonly used by small mammals as roosting and nesting sites (Loeb, 1999), while cavities formed through advanced decay processes may provide shelter for amphibians, reptiles, rodents or mesocarnivores (Dupuis et al., 1995; Duncan, 1999; Bull, 2002; Aubry et al., 2017). Uprooted root plates often create protected nesting sites for ground-nesting birds (De Santo et al., 2003; Wojton & Pitucha, 2020), whereas elevated logs may serve as movement routes or scent-marking locations for carnivores (Trevarrow & Arismendi, 2022; Schwegmann & Storch, 2024). Deadwood also supports extensive epixylic communities of mosses, lichens and fungi, including large fungal fruiting bodies that contribute to both decomposition processes and forest food webs (Humphrey et al., 2002; Runnel et al., 2013; Müller et al., 2015; Brūmelis et al., 2017; Birkemoe et al., 2018). In addition, moisture-retaining nursing logs create favourable sites for seed germination and regeneration of vascular plants, thereby increasing small-scale habitat heterogeneity within forest stands (Faliński, 1978; Szewczyk & Szwagrzyk, 1996; Bače et al., 2012; Staniaszek-Kik et al., 2014).

CWD provides heterogeneous environment that emerge during wood decomposition and long-term exposure to environmental conditions (Maser et al., 1979). The operation of biotic and abiotic factors leads to the formation of different distinct micro-structures occurring on CWD that provide resources or microhabitats essential for particular groups of organisms. Several of these micro-structures correspond conceptually to microhabitats previously described for standing trees under the framework of Tree-related Microhabitats (TreMs; Larrieu et al., 2018), e.g. cavities, bark loss, fungal conks, rot-holes or dendrotelmata, which has become an increasingly important in biodiversity assessment and conservation planning (Martin et al., 2022). Standardized classification of TreMs has substantially improved the integration of biodiversity-related structural attributes into forest inventories and management practices (Kraus et al., 2016b; John et al., 2024; Kadavý et al., 2024).

To date, the features occurring on downed CWD received limited attention compared to structurally analogous TreMs, despite the fact that some of them develop only after tree breakage or uproot. Previous studies of potential structures associated with CWD have focused on selected groups of microhabitats or taxa associated with deadwood, such as fungal communities (Yang et al., 2021), epixylic bryophytes (Müller et al., 2015), hollow logs as den sites (Bull & Heater, 2000) or decay-related substrate conditions (Bače et al., 2012; Kolényová et al., 2024; Lettenmaier et al., 2026). However, a comprehensive framework integrating these structures into a unified system of biologically meaningful deadwood-associated microhabitats is currently lacking. As a result, CWD is being characterized by attributes such as volume, diameter, tree species or decay class, while fine-scale structures occurring on CWD remain not surveyed and their characteristics are rarely used in habitat quality assessments.

Here, we introduce the concept of Deadwood-related Microhabitats (hereafter, DreMs), defined as distinct structures occurring on downed deadwood that may provide shelter, breeding sites, foraging substrate or other critical resources for forest-dwelling organisms. Building upon the conceptual foundations of TreMs, we developed an original classification of DreMs representing biologically meaningful microstructures associated with decomposing CWD in temperate forests. Using an extensive dataset collected in best-preserved old-growth forests representing five major temperate forest types, we quantified the frequency and richness of DreMs and evaluated how these indices are related to CWD characteristics, including species, decomposition class and diameter. By focusing on forests with a high degree of naturalness and long-term ecological continuity, our study provides a reference for CWD-associated structures in temperate forests.

## Methods

### Study area

The study was conducted in five best-preserved old-growth forests in Poland that enjoyed long-term protection. Study sites represented a gradient of forest composition and structural complexity ranging from nearly monospecific coniferous and deciduous stands to highly diverse mixed broadleaved forests (Fig. 1a). The studied forests differed markedly in tree species richness, stand structure and deadwood resources (Przepióra et al., 2026) and comprised:

1. upper montane Norway spruce *Picea abies* forest in the Tatra Mountains (hereafter spruce forest), since 1954 protected as Tatra National Park, dominated by Norway spruce with small admixture of rowan *Sorbus aucuparia* and several other tree species, with mean deadwood volume of 248 m³ ha⁻¹ (Bodziarczyk et al., 2019; Przepióra & Ciach, 2025);
2. montane European beech *Fagus sylvatica* forest in the Bieszczady Mountains (hereafter beech forest), since 1973 protected as Bieszczady National Park and designed as the UNESCO World Heritage Site “Ancient and Primeval Beech Forests of the Carpathians and Other Regions of Europe” (Kirchmeir & Kovarovics, 2020), structurally complex uneven-aged stands with admixtures of sycamore maple *Acer pseudoplatanus*, silver fir *Abies alba* and Norway spruce (Kucharzyk, 2008; Michalik & Szary, 2016), driven mainly by small-scale gap dynamic, with mean deadwood volume of 55 m³ ha⁻¹ (Kacprzyk et al., 2014);
3. willow *Salix* spp.–poplar *Populus* spp. riparian forest in the middle course of the Vistula River valley (hereafter willow–poplar forest), located within Natura 2000 Special Protection Areas and protected within several nature reserves, represents dynamic floodplain ecosystems shaped by periodic inundation, sediment transport and river-channel migration (Gacka- Grzesikiewicz et al., 1995; Przepióra & Ciach, 2022), composed mainly of white willow *Salix alba*, crack willow *Salix fragilis*, white poplar *Populus alba* and black poplar *Populus nigra*, accompanied locally by elm *Ulmus* spp. (Matuszkiewicz, 2001a,b), frequent floods, erosion and ice-related disturbances create heterogeneous age and size structures, contributing to continuous formation of deadwood and habitat diversity;
4. mixed European beech–silver fir forest in the Świętokrzyskie Mountains (hereafter beech–fir forest), since 1950 protected as Świętokrzyski National Park, structurally complex uneven-aged stands, dominated by silver fir and European beech with admixtures of Norway spruce and rowan (Orzechowski & Kędziora, 2023), with mean deadwood volume of 46 m³ ha⁻¹ (Figarski et al., 2014);
5. lowland pedunculate oak *Quercus robur*–small-leaved lime *Tilia cordata*–European hornbeam *Carpinus betulus* forest in Białowieża Forest (hereafter oak–lime–hornbeam forest), since 1932 protected as Białowieża National Park and recognized as a UNESCO World Heritage Site (Kujawa et al., 2016; Jaroszewicz et al., 2019), dominated by European hornbeam, small-leaved lime, Norway spruce and pedunculate oak, accompanied by numerous additional tree species (Faliński, 1986; Keczyński, 2017), have high densities of large and veteran trees, complex vertical structure and mean deadwood volume of 159 m³ ha⁻¹ (Bobiec, 2002).

**Fig. 1.**
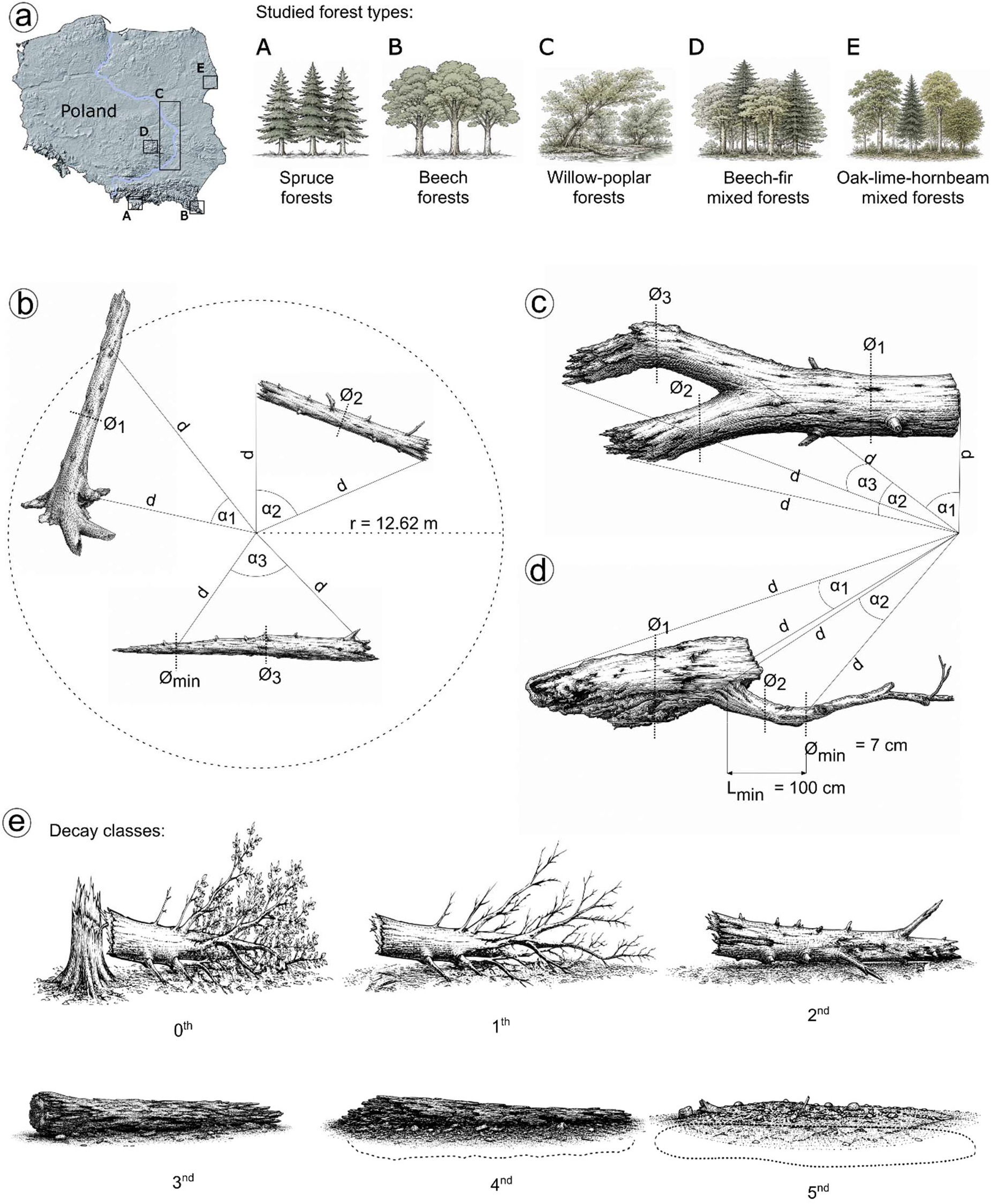
(a) Localization of studied best preserved old-growth forests in Poland: A – Norway spruce *Picea abies*-dominated forests located in Tatra National Park, B – European beech *Fagus sylvatica*-dominated forests located in Bieszczady National Park, C – willow *Salix* spp.-poplar *Populus* spp. riparian forests located in Middle Vistula Valey, D – European beech-silver fir *Abies alba* mixed forests located in Świętokrzyski National Park, D – pedunculate oak *Quercus robus*-small-leaved lime *Tilia cordata*-European hornbeam *Carpinus betulus* mixed forests located in Białowieża National Park; scheme of measurements of downed coarse woody debris (CWD): **(b)** measurement within a circular sample plot, **(c)** measurement of a CWD with forked sections, **(d)** measurement of CWD with sections with an abrupt change in diameter, d – distance from the plot center to the tip of sections, α1, 2, 3 – angle between sighting lines, ø1, 2, 3 – diameter measured at the midpoint of section, ømin – minimum diameter value, Lmin – minimum length, r – radius of the sample plot; **(e)** the stage of decomposition of CWD assessed using a 6-class scale, based on the classification for coniferous deadwood (Maser et al., 1979), which was adapted to the tree species present in the studied forests (Table S1).

In all studied forests, operations such as logging, salvage cutting, planting or deadwood removal are absent or prohibited, allowing forest dynamics to proceed through natural processes and spontaneous disturbances. The study sites are characterized by rich TreM resources, varying between 49 and 61 TreM types per the study site (Przepióra & Ciach, 2023, 2025). Ecological continuity and ongoing natural disturbance agents, including windthrows, bark beetle outbreaks, floods, lightning strikes, fungal decay and the activity of large vertebrates and saproxylic insects contribute to high quantity and diversity of CWD (Jaroszewicz et al., 2019; Przepióra & Ciach, 2022).

### Field methods

The fieldwork was conducted on 423 circular study plots (Tatra Mts., N = 90; Bieszczady Mts., N = 109; Middle Vistula Valley, N = 91; Świętokrzyskie Mts., N = 30; Białowieża Forest, N = 103), each with a radius of 12.62 m (0.05 ha; Fig. 1b). Although most study plots being located in national parks or reserves, selected forest patches had to display a high degree of naturalness, with no visible signs of human interference, such as tree removal, tree planting or cattle grazing. A minimum distance between plots in each of studied forests was maintained at more than twice the stand height, with an average distance between adjacent plots of 152.9 m ±81.2 SD (range 41.0–466.4 m). To ensure the visibility of CWD, fieldwork was carried out during the leafless season, either before the development or after the decay of forest floor vegetation, and without snow cover, from March 2019 to October 2024.

On each plot, CWD was surveyed, ranging from individual deadwood pieces, i.e. branches, piece of trunk to entire fallen trees composed of multiple interconnected trunk and branch sections. CWD were included when they met the minimum size criteria of ≥ 7 cm in diameter at the thinner end and ≥ 100 cm in length (Fig. 1b). For fallen trees, in particular large trees with complex architecture composed of multiple interconnected trunk and branch sections, all eligible sections were measured separately; however, they were treated collectively as single debris. A section was defined as a continuous woody segment meeting the minimum size criteria, extending either between successive branching points (Fig. 1c) or between a branching point and a point where the stem diameter changed abruptly (Fig. 1d). For each CWD lying within the study plot limit, the position of each section was recorded by measuring the azimuth and distance of both ends from the plot centre and the diameter was measured at the midpoint of its length. The species of each debris was identified, while taxa with similar wood/bark features, such as crack willow and white willow, white poplar and black poplar, and elms (wych elm *Ulmus glabra*, European white elm *Ulmus laevis* and field elm *Ulmus minor*) were attributed to genus.

The stage of decomposition of CWD was assessed using a 6-class scale, based on the classification for coniferous deadwood (Maser et al., 1979), which was adapted to the tree species present in the studied forests (Table S1; Fig. 1e). Dominant decay class was assigned for each CWD (hereafter, main decay class). As fragments of a given CWD may differ in decomposition stage and lead to occurrence of multiple decay classes on single CWD, the presence of additional decay class was assigned if its fragment met the criteria of CWD, i.e. diameter ≥ 7 cm and length ≥ 100 cm.

DreMs present on CWD were inventoried using a original catalogue (Appendix 1). The list of groups and types of microhabitats was adapted from those identified for standing trees (Kraus et al., 2016a; Larrieu et al., 2018). Based on field reconnaissance, microhabitats from the catalogue for standing trees were selected if they also were found on lying deadwood, while other have been included when they had documented evidence for use by organisms for development, reproduction, foraging or shelter (Appendix 2). Polypores and other fungal fruiting bodies serve as habitats for specialized invertebrates (Birkemoe et al., 2018) and the composition of fungivorous insect assemblages varies with the degree of fruiting body decomposition (Jonsen & Nordlander, 2004). Cracks and spaces resulting from the decomposition of CWD core may be used as shelter or breeding sites by various vertebrates and invertebrates (Carey & Johnson, 1995; Loeb, 1999; Fritts et al., 2015; Zumr et al., 2024). Cavities are important microhabitats for numerous organisms, serving as breeding sites for birds and providing suitable developmental conditions for saproxylic insects (van der Hoek et al., 2017; Zumr et al., 2024). While birds primarily use cavities in standing trees, those in lying deadwood can provide nesting sites for facultative cavity-nesters (De Santo et al., 2003; Martin et al., 2004; Karpińska et al., 2022). Cavities were classified with distinctions made between small (≤ 10 cm) and large (> 10 cm) cavities formed by wood decay (saproxylogenic) and those excavated by woodpeckers (woodpecker breeding cavities). Woodpecker foraging holes are used by thrushes, e.g. common blackbird *Turdus merula* for nest placement (Tomiałojć, 1993). Loose, peeling bark serves as a habitat for invertebrates (Zumr et al., 2024). Vegetation covering the surface of CWD supports diverse assemblages of invertebrates, with different vegetation types creating distinct structural and microclimatic conditions that support different arthropod communities (Stokland et al., 2012; Zumr et al., 2024; Porto et al., 2025). Many insects are associated with bird or mammal nests (Jaworski et al., 2022), while stagnant water is microhabitat for amphibians and insects (Gossner & Petermann, 2022). Larval galleries, tunnels and bore holes created by xylophagous insects are reused by a range of saproxylic organisms (Marquis & Lill, 2007) and large emerging holes (> 2 cm in diameter) provide shelter or refugia for small vertebrates, including lizards and snakes (Gottfried et al., 2019; Borczyk et al., 2022). Uprooted trees with exposed root plates create pit-and-mound microtopography that increases habitat heterogeneity and provide shelter and nesting sites for birds associated with ground and shrub layers (Wesołowski, 1983; Tomiałojć, 1993; Ulanova, 2000; Wojton & Pitucha, 2020). Two types of uprooted trees were inventoried— with soil or without soil, with ≥ 0.01 m³ of soil required for classification. Accumulations ≥ 0.01 m³ of organic matter within CWD, such as decomposing leaves and small branches, retain higher moisture levels for extended periods, providing shelter for moisture-dependent organisms like amphibians (DeMaynadier & Hunter, 1995). These accumulations also serve as substrate for plants that would otherwise not thrive solely on decomposing wood. Charred and partially burned wood ≥ 0.1 m², which serves as a substrate for pyrophilous insects and fungi (Penttilä & Kotiranta, 1996; Gutowski et al., 2020).

The inventory of DreMs included 14 groups divided into 30 types (Table 2). Cataloguing involved confirming the presence or absence of each DreM on a given CWD; if a given DreM was present on more than one sections of the CWD (e.g. on both the trunk and branches of a fallen tree), it was recorded only once. To minimize potential observer bias known for TreMs inventories (Paillet et al., 2015), DreMs were recorded by a single observer (FP). Only the visible parts of the CWD were examined and the surface in contact with the ground was not inspected. Each of debris was scrutinized for a minimum of 3 min. CWD came from spruce forests (N = 1,624), beech forests (N = 1,542), followed by oak-lime-hornbeam forests (N = 1,409), willow-poplar forests (N = 1,068) and beech-fir forests (N = 360) (Table S2).

**Table 1.** Characteristics of downed coarse woody debris (CWD) and Deadwood-related Microhabitats (DreMs) found in old-growth forests in Poland. Mean ±SD and range of **(a)** CWD diameter, **(b)** CWD volume, **(c)** number of sections i.e. number of interconnected trunks and branches making up a CWD and **(d)** DreM richness (number of DreM types found on and single debris) calculated for taxa with N ≥ 30 (see Table S2 for number of surveyed CWD): all CWD pooled (total), deciduous taxa, coniferous taxa, Norway spruce *Picea abies*, European beech *Fagus sylvatica*, crack willow *Salix fragilis* and white willow *Salix alba* pooled, European hornbeam *Carpinus betulus*, white poplar *Populus alba* and black poplar *Populus nigra* pooled, silver fir *Abies alba*, small-leaved lime *Tilia cordata*, rowan *Sorbus aucuparia*, pedunculate oak *Quercus robur*, goat willow *Salix caprea*, European aspen *Populus tremula*, European ash *Fraxinus excelsior* and silver birch *Betula pendula*.

| Category | Mean | SD | Range |
| --- | --- | --- | --- |
| **(a) CWD diameter** |  |  |  |
| Total | 18.0 | 10.7 | 7.0-100.0 |
| Deciduous | 17.5 | 11.4 | 7.0-100.0 |
| Coniferous | 18.9 | 9.5 | 7.0-75.0 |
| Norway spruce | 18.9 | 9.4 | 7.0-75.0 |
| European beech | 19.3 | 13.5 | 7.0-100.0 |
| Crack willow and white willow | 15.6 | 7.6 | 7.0-52.0 |
| European hornbeam | 20.6 | 12.1 | 7.0-70.0 |
| White poplar and black poplar | 17.6 | 8.9 | 7.0-62.0 |
| Silver fir | 18.9 | 10.3 | 7.0-50.5 |
| Small-leaved lime | 13.3 | 7.9 | 7.0-54.5 |
| Rowan | 14.2 | 5.1 | 7.0-28.0 |
| Pedunculate oak | 18.1 | 14.5 | 7.0-65.0 |
| Goat willow | 15.2 | 5.3 | 7.0-31.0 |
| European aspen | 19.2 | 11.6 | 7.0-65.0 |
| European ash | 20.0 | 9.4 | 8.0-45.7 |
| Silver birch | 14.5 | 5.1 | 7.0-27.0 |
| **(b) CWD volume** |  |  |  |
| Total | 0.2 | 0.5 | 0.004-6.6 |
| Deciduous | 0.2 | 0.5 | 0.004-6.6 |
| Coniferous | 0.3 | 0.4 | 0.004-5.4 |
| Norway spruce | 0.3 | 0.4 | 0.004-5.4 |
| European beech | 0.3 | 0.6 | 0.004-5.8 |
| Crack willow and white willow | 0.2 | 0.3 | 0.005-2.8 |
| European hornbeam | 0.3 | 0.5 | 0.005-3.9 |
| White poplar and black poplar | 0.3 | 0.4 | 0.008-3.6 |
| Silver fir | 0.3 | 0.5 | 0.005-4.2 |
| Small-leaved lime | 0.1 | 0.2 | 0.004-1.0 |
| Rowan | 0.1 | 0.1 | 0.007-0.6 |
| Pedunculate oak | 0.4 | 1.1 | 0.004-5.2 |
| Goat willow | 0.1 | 0.1 | 0.009-0.6 |
| European aspen | 0.3 | 0.5 | 0.005-3.0 |
| European ash | 0.4 | 0.5 | 0.008-1.9 |
| Silver birch | 0.1 | 0.2 | 0.005-0.8 |
| **(c) Number of sections** |  |  |  |
| Total | 1.1 | 0.7 | 1.0-18.0 |
| Deciduous | 1.2 | 0.9 | 1.0-18.0 |
| Coniferous | 1.0 | 0.1 | 1.0-3.0 |
| Norway spruce | 1.0 | 0.1 | 1.0-3.0 |
| European beech | 1.2 | 0.8 | 1.0-13.0 |
| Crack willow and white willow | 1.1 | 0.6 | 1.0-8.0 |
| European hornbeam | 1.5 | 1.7 | 1.0-18.0 |
| White poplar and black poplar | 1.1 | 0.5 | 1.0-6.0 |
| Silver fir | 1.0 | 0.0 | 1.0-1.0 |
| Small-leaved lime | 1.0 | 0.3 | 1.0-3.0 |
| Rowan | 1.3 | 0.7 | 1.0-4.0 |
| Pedunculate oak | 1.8 | 2.2 | 1.0-14.0 |
| Goat willow | 1.2 | 0.7 | 1.0-5.0 |
| European aspen | 1.5 | 1.7 | 1.0-11.0 |
| European ash | 1.4 | 0.9 | 1.0-5.0 |
| Silver birch | 1.1 | 0.4 | 1.0-3.0 |
| **(d) DreM richness** |  |  |  |
| Total | 2.1 | 1.6 | 0.0-15.0 |
| Deciduous | 2.2 | 1.8 | 0.0-15.0 |
| Coniferous | 2.0 | 1.4 | 0.0-13.0 |
| Norway spruce | 1.9 | 1.3 | 0.0-8.0 |
| European beech | 2.1 | 1.9 | 0.0-14.0 |
| Crack willow and white willow | 2.8 | 1.7 | 0.0-11.0 |
| European hornbeam | 2.7 | 2.0 | 0.0-15.0 |
| White poplar and black poplar | 2.7 | 2.0 | 0.0-13.0 |
| Silver fir | 2.7 | 2.0 | 0.0-13.0 |
| Small-leaved lime | 1.7 | 1.2 | 0.0-6.0 |
| Rowan | 2.8 | 1.6 | 0.0-8.0 |
| Pedunculate oak | 2.1 | 1.5 | 0.0-7.0 |
| Goat willow | 2.5 | 1.3 | 0.0-7.0 |
| European aspen | 2.1 | 1.9 | 0.0-7.0 |
| European ash | 2.8 | 1.7 | 0.0-6.0 |
| Silver birch | 3.6 | 2.4 | 0.0-12.0 |

**Table 2.**
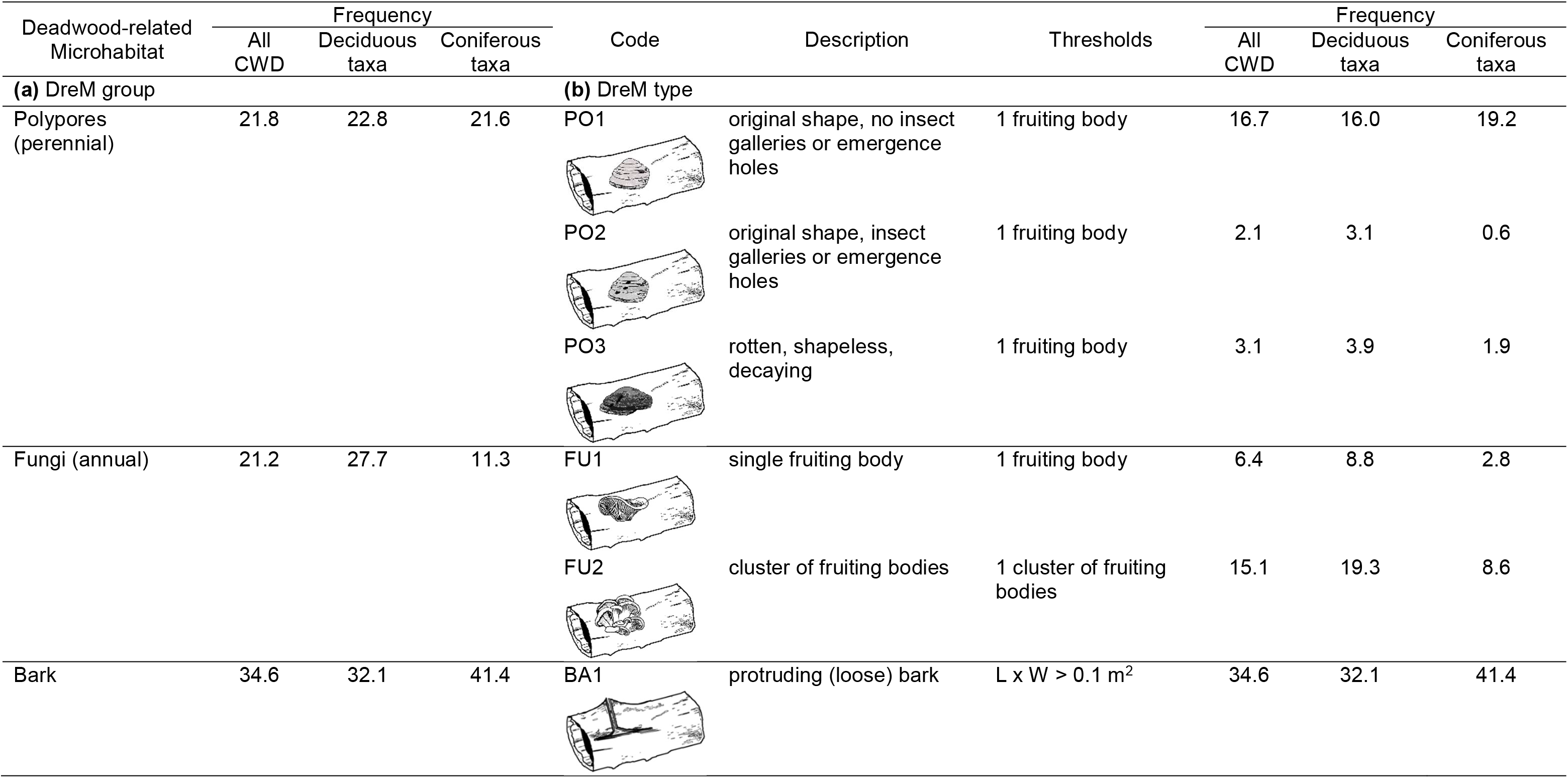
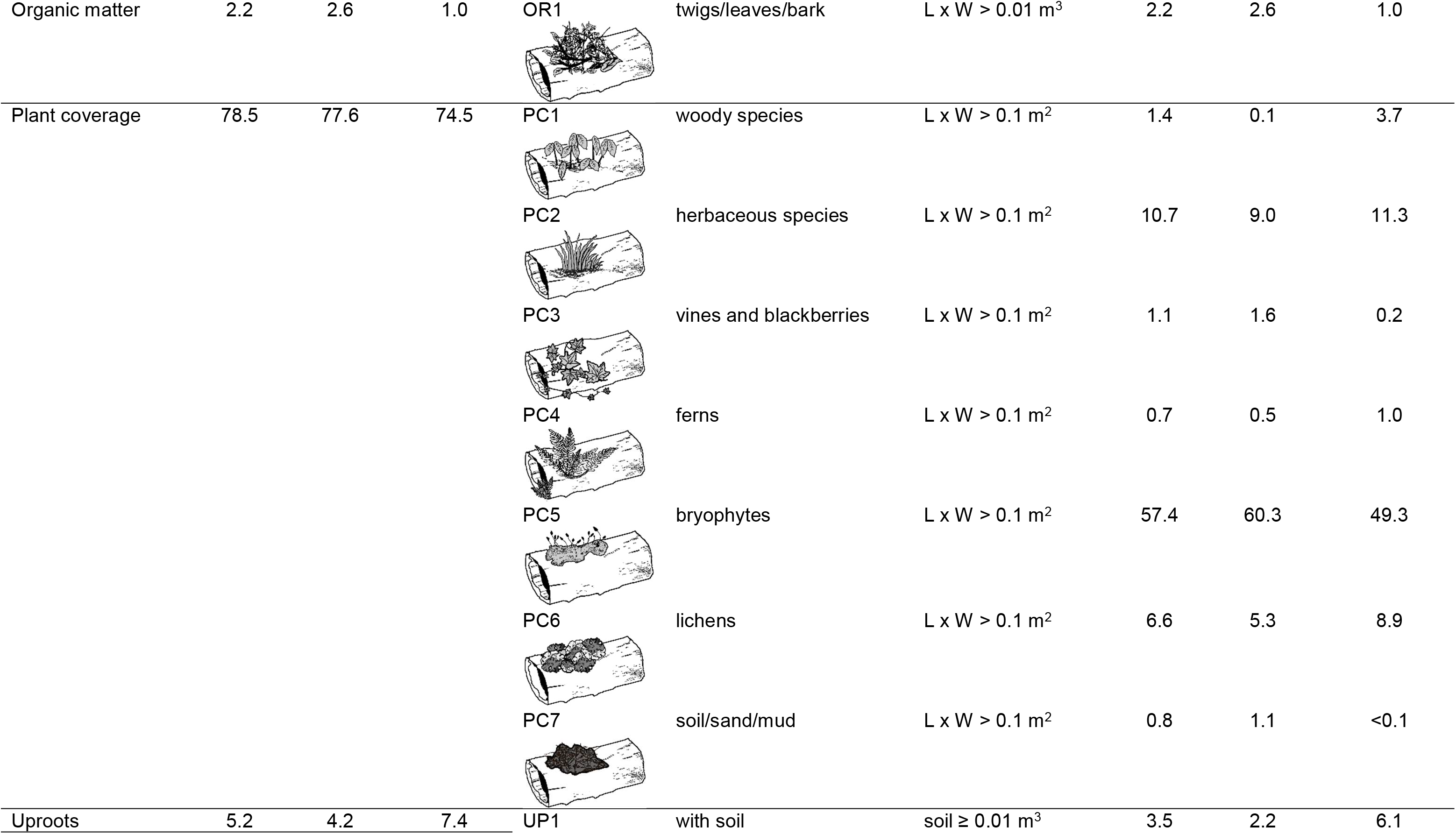
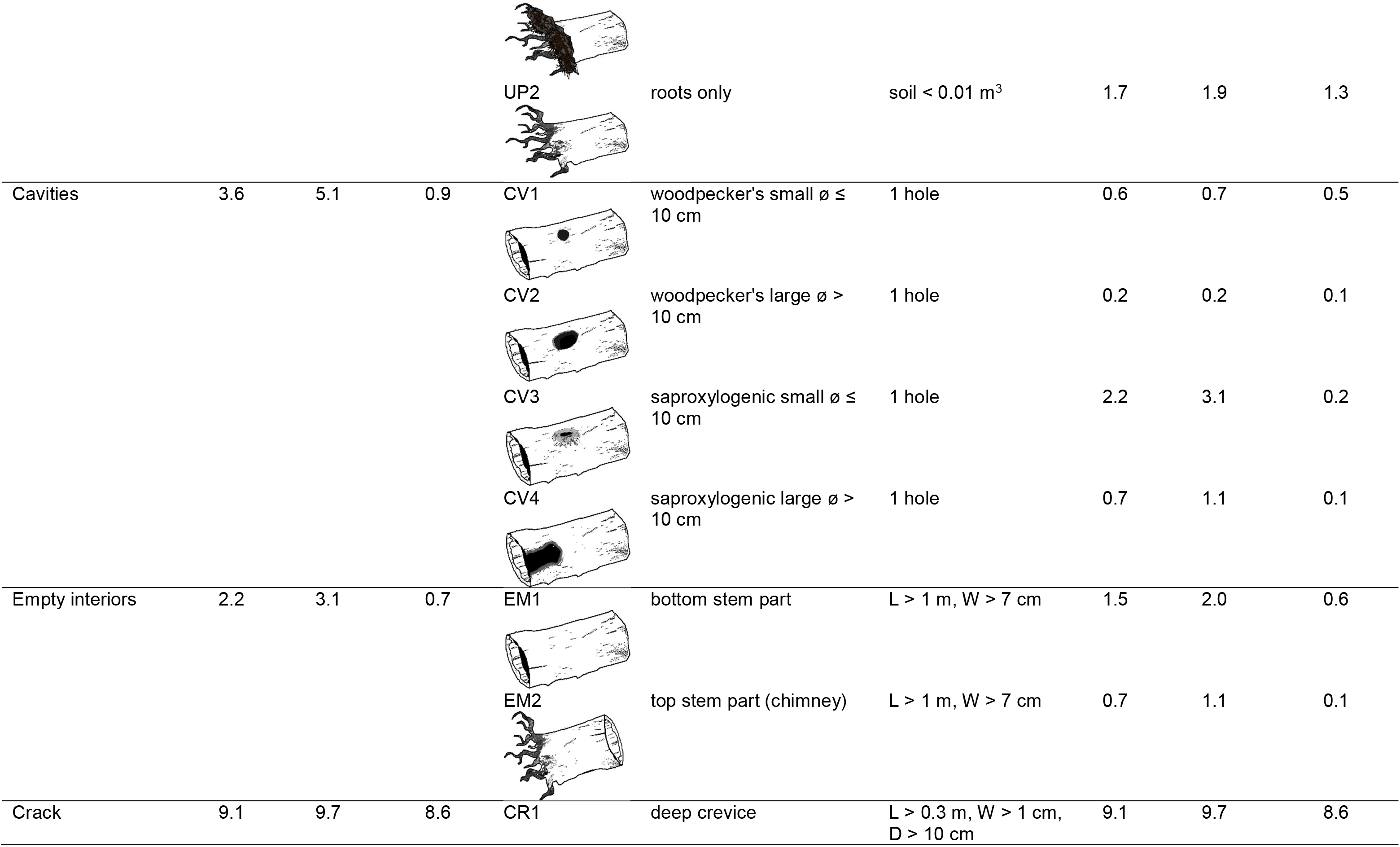
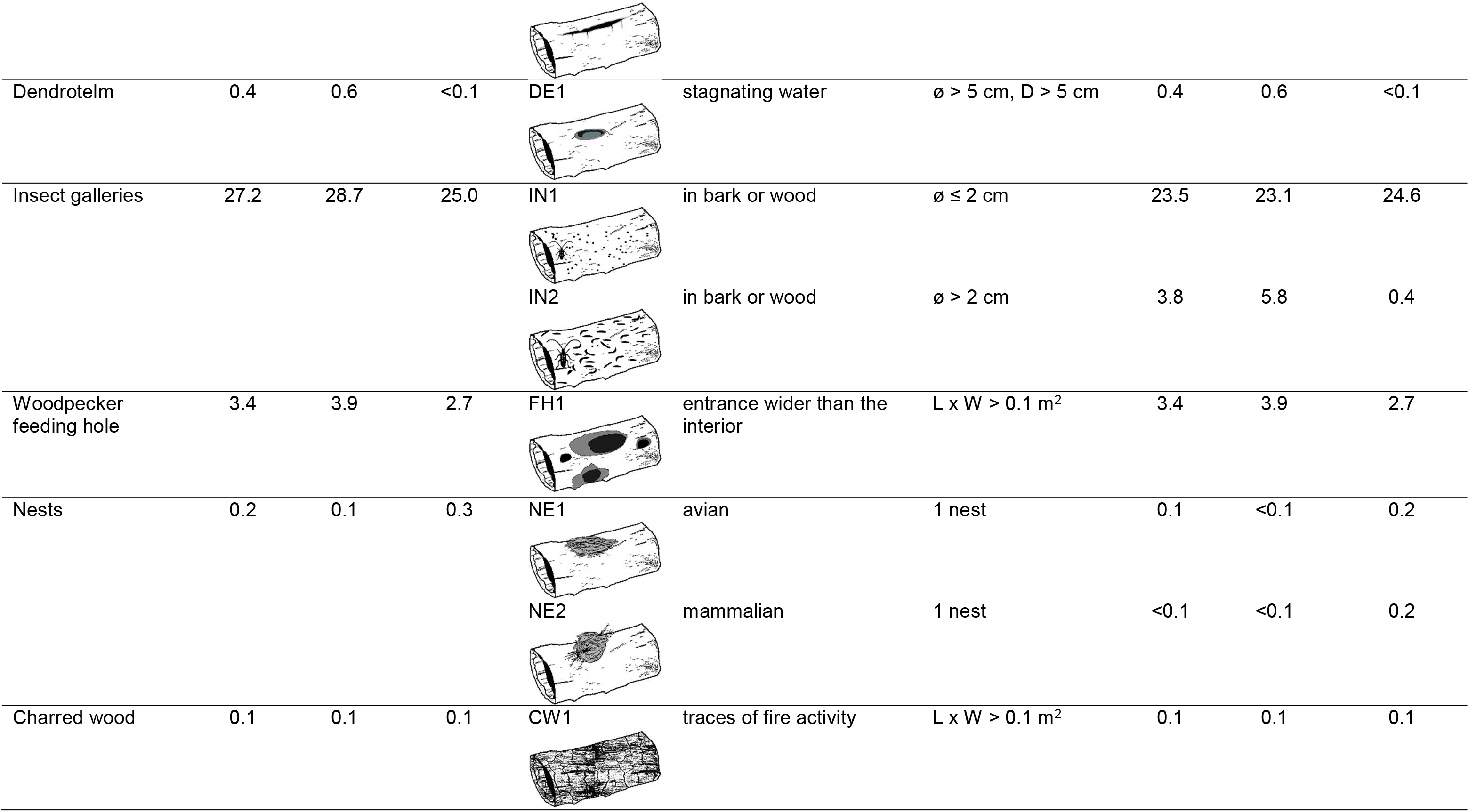
Frequency (percentage) of downed coarse woody debris (CWD) bearing specific Deadwood-related Microhabitats (DreMs) **(a)** group or **(b)** type calculated for all recorded CWD (N = 6003) and deciduous (N = 3702) or coniferous taxa (N = 2107) found in old-growth forests in Poland. L – length, W – width, D – depth, ø – diameter, see Appendix 1 for examples of associated organisms.

| Deadwood-related Microhabitat | Frequency |  |  | Code | Description | Thresholds | Frequency |  |  |
| --- | --- | --- | --- | --- | --- | --- | --- | --- | --- |
|  | All CWD | Deciduous taxa | Coniferous taxa |  |  |  | All CWD | Deciduous taxa | Coniferous taxa |
| **(a) DreM group** |  |  |  | **(b) DreM type** |  |  |  |  |  |
| Polypores (perennial) | 21.8 | 22.8 | 21.6 | PO1 | original shape, no insect galleries or emergence holes | 1 fruiting body | 16.7 | 16.0 | 19.2 |
|  |  |  |  | PO2 | original shape, insect galleries or emergence holes | 1 fruiting body | 2.1 | 3.1 | 0.6 |
|  |  |  |  | PO3 | rotten, shapeless, decaying | 1 fruiting body | 3.1 | 3.9 | 1.9 |
| Fungi (annual) | 21.2 | 27.7 | 11.3 | FU1 | single fruiting body | 1 fruiting body | 6.4 | 8.8 | 2.8 |
|  |  |  |  | FU2 | cluster of fruiting bodies | 1 cluster of fruiting bodies | 15.1 | 19.3 | 8.6 |
| Bark | 34.6 | 32.1 | 41.4 | BA1 | protruding (loose) bark | L x W > 0.1 m<sup>2</sup> | 34.6 | 32.1 | 41.4 |
| Organic matter | 2.2 | 2.6 | 1.0 | OR1 | twigs/leaves/bark | $L \times W > 0.01 \text{ m}^3$ | 2.2 | 2.6 | 1.0 |
| Plant coverage | 78.5 | 77.6 | 74.5 | PC1 | woody species | $L \times W > 0.1 \text{ m}^2$ | 1.4 | 0.1 | 3.7 |
| | | | | PC2 | herbaceous species | $L \times W > 0.1 \text{ m}^2$ | 10.7 | 9.0 | 11.3 |
| | | | | PC3 | vines and blackberries | $L \times W > 0.1 \text{ m}^2$ | 1.1 | 1.6 | 0.2 |
| | | | | PC4 | ferns | $L \times W > 0.1 \text{ m}^2$ | 0.7 | 0.5 | 1.0 |
| | | | | PC5 | bryophytes | $L \times W > 0.1 \text{ m}^2$ | 57.4 | 60.3 | 49.3 |
| | | | | PC6 | lichens | $L \times W > 0.1 \text{ m}^2$ | 6.6 | 5.3 | 8.9 |
| | | | | PC7 | soil/sand/mud | $L \times W > 0.1 \text{ m}^2$ | 0.8 | 1.1 | <0.1 |
| Uproots | 5.2 | 4.2 | 7.4 | UP1 | with soil | $\text{soil} \geq 0.01 \text{ m}^3$ | 3.5 | 2.2 | 6.1 |

|  |  |  |  |  |  |  |  |  |  |  |
| --- | --- | --- | --- | --- | --- | --- | --- | --- | --- | --- |
|  |  |  |  | UP2 | roots only | soil < 0.01 m<sup>3</sup> | 1.7 | 1.9 | 1.3 |  |
| Cavities | 3.6 | 5.1 | 0.9 | CV1 | woodpecker's small $\varnothing \leq 10$ cm | 1 hole | 0.6 | 0.7 | 0.5 | |
| | | | | CV2 | woodpecker's large $\varnothing > 10$ cm | 1 hole | 0.2 | 0.2 | 0.1 | |
| | | | | CV3 | saproxylogenic small $\varnothing \leq 10$ cm | 1 hole | 2.2 | 3.1 | 0.2 | |
| | | | | CV4 | saproxylogenic large $\varnothing > 10$ cm | 1 hole | 0.7 | 1.1 | 0.1 | |
| Empty interiors | 2.2 | 3.1 | 0.7 | EM1 | bottom stem part | L > 1 m, W > 7 cm | 1.5 | 2.0 | 0.6 |  |
|  |  |  |  | EM2 | top stem part (chimney) | L > 1 m, W > 7 cm | 0.7 | 1.1 | 0.1 |  |
| Crack | 9.1 | 9.7 | 8.6 | CR1 | deep crevice | L > 0.3 m, W > 1 cm,<br>D > 10 cm | 9.1 | 9.7 | 8.6 |  |
| Dendrotelm | 0.4 | 0.6 | <0.1 | DE1 | stagnating water | $\varnothing > 5 \text{ cm}, D > 5 \text{ cm}$ | 0.4 | 0.6 | <0.1 |
| Insect galleries | 27.2 | 28.7 | 25.0 | IN1 | in bark or wood | $\varnothing \leq 2 \text{ cm}$ | 23.5 | 23.1 | 24.6 |
| | | | | IN2 | in bark or wood | $\varnothing > 2 \text{ cm}$ | 3.8 | 5.8 | 0.4 |
| Woodpecker feeding hole | 3.4 | 3.9 | 2.7 | FH1 | entrance wider than the interior | $L \times W > 0.1 \text{ m}^2$ | 3.4 | 3.9 | 2.7 |
| Nests | 0.2 | 0.1 | 0.3 | NE1 | avian | 1 nest | 0.1 | <0.1 | 0.2 |
|  |  |  |  | NE2 | mammalian | 1 nest | <0.1 | <0.1 | 0.2 |
| Charred wood | 0.1 | 0.1 | 0.1 | CW1 | traces of fire activity | $L \times W > 0.1 \text{ m}^2$ | 0.1 | 0.1 | 0.1 |

### Data handling and analyses

For each CWD, its volume (m³) was calculated using the formula for the midsection:

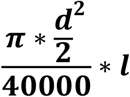

where *d* represents the diameter (cm) at the midpoint of the section, and *l* is the length of the section (m). The diameter of the thickest measured section (hereafter, CWD diameter), the total volume of all sections composing the CWD (hereafter, CWD volume), the total number of sections, and the total number of DreM types present on the entire CWD (hereafter, DreM richness) were calculated.

Mean, standard deviation (SD) and range of CWD diameter, CWD volume, DreM richness and number of sections of making up a CWD and percentage of CWD in main decay class were calculated for (1) all CWD pooled, (2) deciduous and coniferous tree taxa separately and (3) each tree taxa with N ≥ 30. Differences in CWD diameter, CWD volume, number of sections and DreM richness between deciduous and coniferous tree taxa were tested using Student t-test, whereas differences in the distribution of main decay classes between deciduous and coniferous taxa were assessed using Pearson’s chi-squared test. The frequency, i.e. the percentage of CWD bearing specific DreM group and types and the percentage of CWD in a given decay class that bear additional wood decay class were calculated for all CWD pooled deciduous and coniferous tree taxa separately and each tree taxa with N ≥ 30.

The relationships between the CWD characteristics and DreM richness were analyzed using Generalized Linear Mixed Models with a Conway-Maxwell Poisson error distribution and a log link function. As the number of unidentified CWD was low (3.2%), the data was limited to deciduous or coniferous taxa (N = 5809). Prior to the modelling procedures, the explanatory variables were tested for co-linearity using Pearson’s correlation. CWD volume was excluded from the set of explanatory variables as it was correlated with CWD diameter (r = 0.76, p < 0.001). Continuous explanatory variables included CWD diameter and number of sections, while categorical variables included taxa (deciduous vs coniferous), presence of additional decay class (presence vs absence) and main decay classes (set as dummy variables: 1 if the CWD is in a given main decay class or 0 if the CWD is not in a given main decay class). The identification numbers of the study plot (423 levels) and study area (5 levels) were added as random effects to the model to counteract the overdispersion and zero-inflation of residuals (Kassahun et al., 2014).

The Akaike Information Criterion corrected for small sample size (AICc) of the model with random effects were lower from the model without random effects (ΔAICc = 628.9) and null model, i.e. model that contain only an intercept (ΔAICc = 2744.1). The residuals of the final model exhibited no dispersion (dispersion = 0.98; p = 0.952; simulation-based dispersion tests) and zero-inflation (ratioObsSim = 1.22; p = 0.272). The differences in mean DreM richness predicted by the model between tree taxa and CWD in different decay classes were tested using post-hoc Tukey test. All analyses were conducted in R 3.5.0 (CoreTeam R., 2017). Models, their diagnostics and post-hoc Tukey test were performed with the glmmTMB (Magnusson et al., 2017), DHARMa (Hartig and Hartig, 2017) and emmeans (Lenth et al. 2018) packages, respectively.

## Results

A total of 6,003 CWD were examined, with 61.7% representing deciduous, 35.1% coniferous taxa. Majority of deciduous debris (76.9%) was identified to the species or genus, while all coniferous debris was identified to species. Deciduous taxa had lower mean CWD diameter (t = 5.20, p < 0.001) (Table 1a) and lower mean CWD volume (t = 3.35, p < 0.001) (Table 1b), while had a higher number of sections (t = -11.27, p < 0.001) (Table 1c) and greater mean DreM richness (t = -3.97, p < 0.001)compared to coniferous (Table 1d). Deciduous and coniferous CWD differed in their main decay-class distribution (χ² = 173.5, df = 5, p < 0.001): coniferous taxa were more frequently represented by 2^nd^ decay class, whereas deciduous by 3^rd^-5^th^ decay classes (Table S3).

All DreM groups (Table 2a) and DreM types (Table 2b) were found on both deciduous and coniferous taxa. The most common DreM groups with frequency > 20% across all CWD were plant cover, bark, insect galleries, polypores and fungi, whereas all remaining DreM groups occurred with frequency < 10% (Table 2a). The most frequent DreM groups on all CWD were also the most frequent on both deciduous and coniferous taxa (Table 2a). DreM types with the highest frequency reaching > 20% on all CWD, as well as on both deciduous and coniferous taxa, were patches of bryophytes (PC5), loose bark patches (BA1) and small insect galleries (IN1) (Table 2b). These DreM types were also the most frequent on Norway spruce, crack and white willow, white and black poplar, silver fir, rowan, goat willow *Salix caprea* and European aspen *Populus tremula* (Fig. 2).

**Fig. 2.**
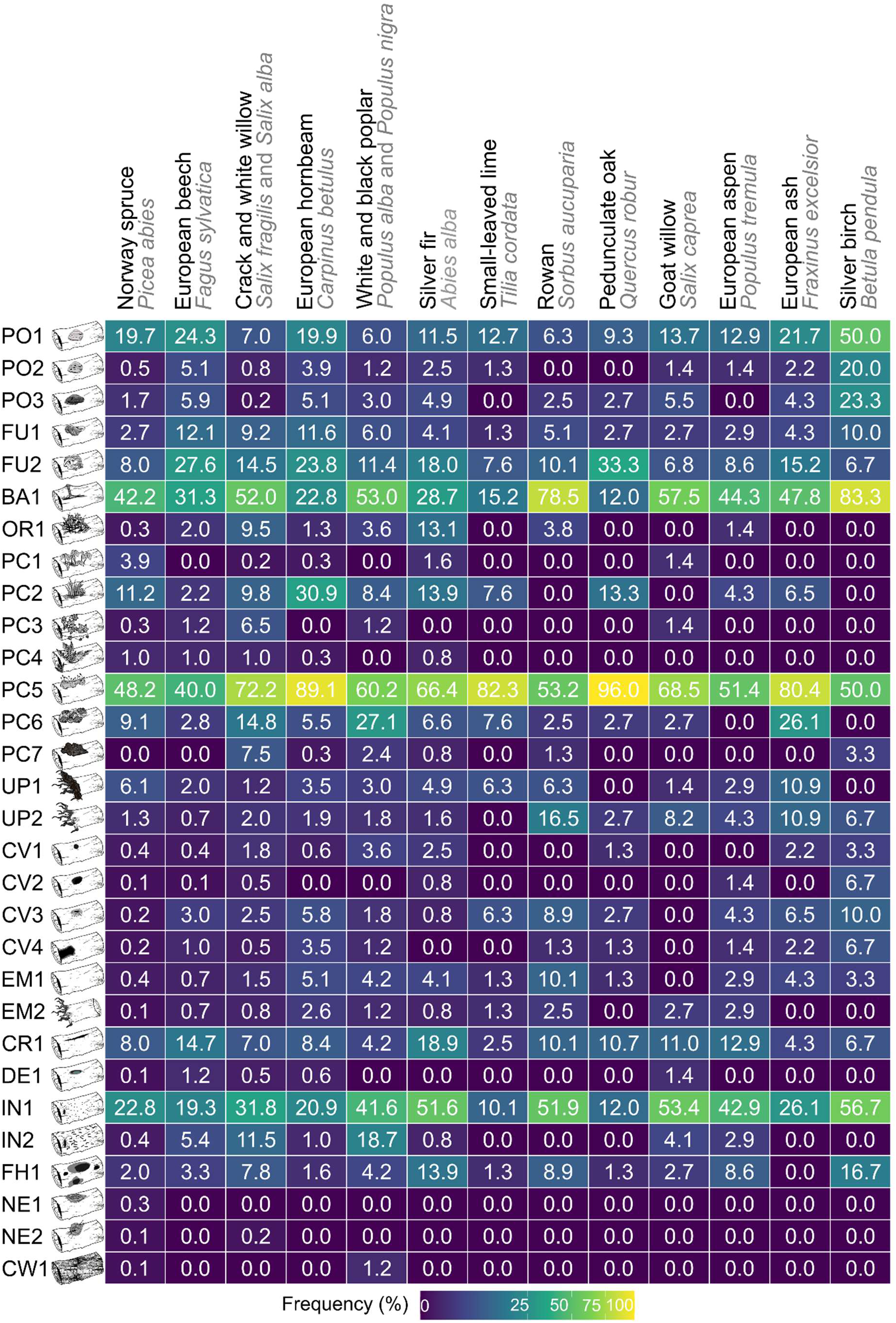
Frequency of occurrence of Deadwood-related Microhabitat (DreMs) (percentage of downed coarse woody debris (CWD) bearing specific type of DreMs) found in old-growth forests in Poland. Calculations included taxa with N ≥ 30: Norway spruce *Picea abies*, European beech *Fagus sylvatica*, crack willow *Salix fragilis* and white willow *Salix alba* pooled, European hornbeam *Carpinus betulus*, white poplar *Populus alba* and black poplar *Populus nigra* pooled, silver fir *Abies alba*, small-leaved lime *Tilia cordata*, rowan *Sorbus aucuparia*, pedunculate oak *Quercus robur*, goat willow *Salix caprea*, European aspen *Populus tremula*, European ash *Fraxinus excelsior* and silver birch *Betula pendula*; see Table 2 and Table S2 for descriptions of DreM codes and number of surveyed CWD, respectively.

The highest number of DreM types was recorded on CWD in the 2^nd^ decay class (30 types) and the 3^rd^ and 4^th^ decay classes (29 types each). Fewer DreM types were recorded in the 1^st^ decay class (26 types), whereas the lowest numbers were observed in the 5^th^ and 0^th^ decay classes – 20 and 19 types, respectively (Fig. 3). Patches of bryophytes (PC5) were the dominant DreM type, being the most frequent in the 0^th^, 1^st^, and 3^rd^–5^th^ decay classes, and the second most frequent in the 2^nd^ decay class on which protruding bark (BA1) was the dominant. The second most frequent DreM varied among decay classes: patches of lichens (PC6) was ranked the second in the 0^th^ decay class, protruding bark (BA1) in the 1^st^, insect galleries (IN1) in the 3^rd^ and patches of herbaceous plant cover (PC2) in the 4^th^ and 5^th^ decay classes (Fig. 3). Among DreM types occurring at frequencies < 20%, fruiting bodies of perennial polypores (PO1) and annual fungi (FU2) changes in frequency along the wood decomposition gradient, reaching their highest frequencies in the 2^nd^ and 1^st^ decay classes, respectively (Fig. 3), while remaining DreM types in this frequency range showed no consistent pattern across decay classes (Fig. 3). Among CWD containing more than one decay class, the 3^rd^ and 4^th^ decay classes as additional class were the most frequently recorded on all CWD, as well as on both deciduous and coniferous taxa (Table S4).

**Fig. 3.**
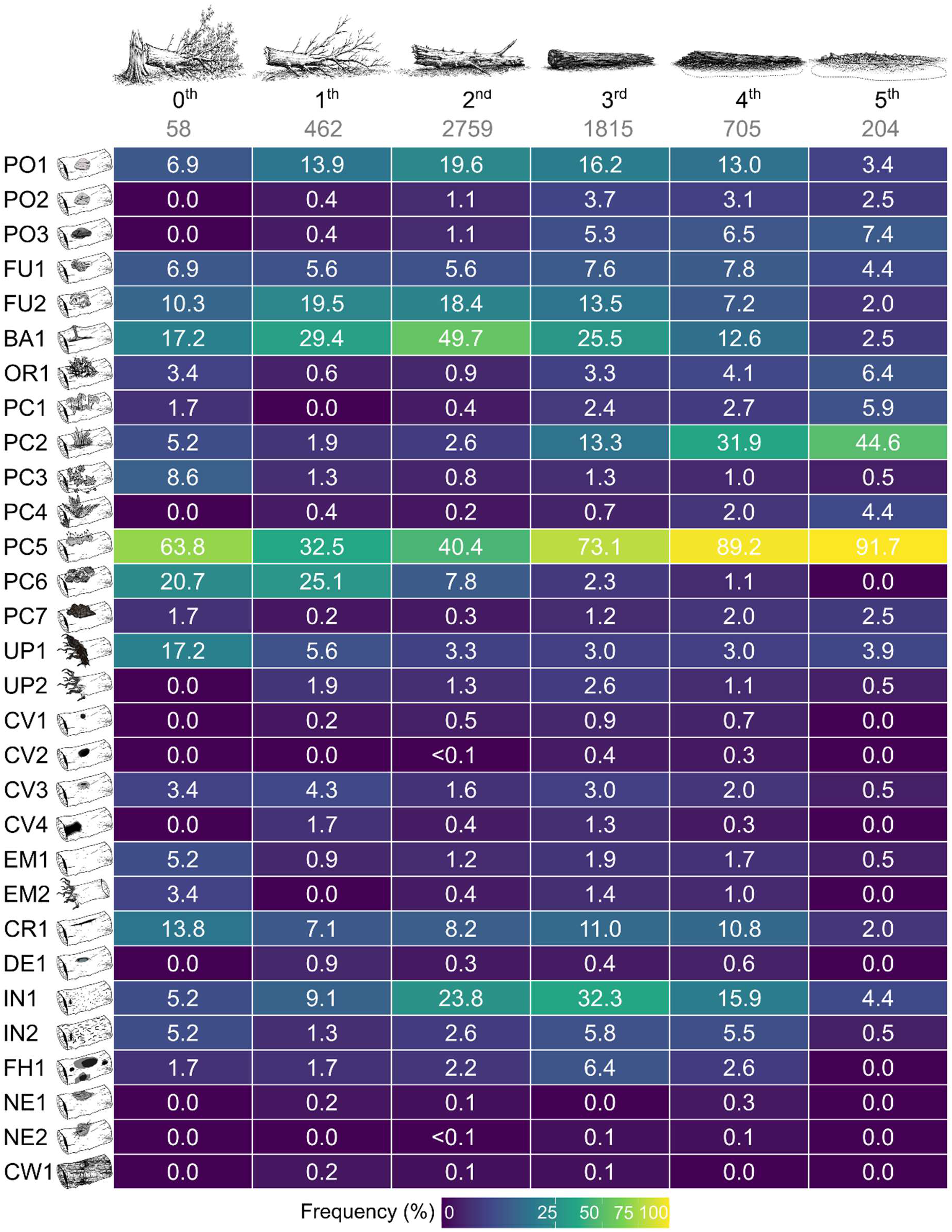
Frequency of occurrence of Deadwood-related Microhabitat (DreMs) (percentage of downed coarse woody debris (CWD) bearing specific type of DreMs) calculated for six main class of decay found in old-growth forests in Poland. Sample sizes are reported below each decay class; see Table 2 and Table S1 for descriptions of DreM codes and decay classes classification, respectively.

DreM richness was correlated with CWD diameter, number of sections, taxa, main decay class and the presence of additional decay class (Table 3). The predicted DreM richness was higher in deciduous than coniferous taxa (Fig. 4a) and increased exponentially with the CWD diameter (Fig. 4b-c). In CWD with diameter > 60 cm for every 10 cm increase in diameter, the number of predicted DreM increased by approximately 30% on both deciduous (Fig. 4b) and coniferous taxa (Fig. 4c). The predicted DreM richness was elevated in the intermediate stages of decomposition, reaching the highest values in decay classes 2^nd^, 3^rd^ and 4^th^ decay classes (Fig. 4d). Although the predicted DreM richness was highest in the 0^th^ decay class, it did not differ from other decay classes. In contrast, CWD in 1^st^ and 5^th^ decay classes has lower DreM richness than CWD in the 2^nd^–4^th^ decay classes (Fig. 4d). The presence of additional decay classes was associated with the increase in the DreM richness (Table 3).

**Fig. 4.**
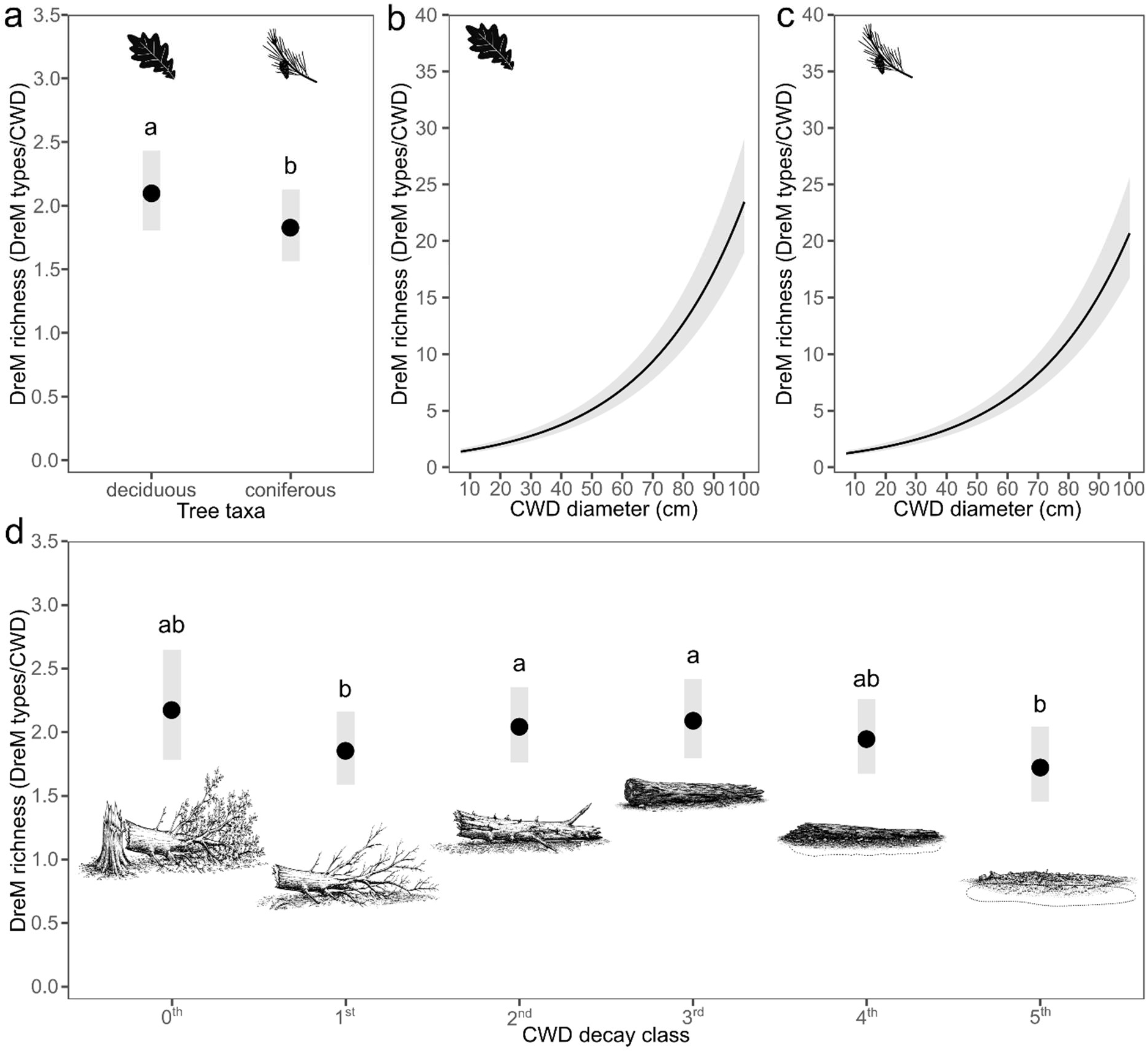
Deadwood-related Microhabitat (DreM) richness (total number of DreM types recorded on downed coarse woody debris (CWD)) found in old-growth forests in Poland. Differences in DreM richness between **(a)** deciduous and coniferous CWD; the relationships between DreM richness and CWD diameter of **(b)** deciduous or **(c)** coniferous taxa and **(d)** DreM richness found on CWD in main classes of decay. Means – filled black circles (a, d) or black lines (b, c), and confidence intervals – grey ribbons (a, b, c, d) are products of the generalized linear-mixed models (see Table 3). Differences among groups were assessed using ANOVA with Tukey’s post hoc tests; different letters (a–b) indicate differences significant at p < 0.05.

**Table 3.** Model describing the relationships between Deadwood-related Microhabitats (DreMs) and characteristics of downed coarse woody debris (CWD) found in old-growth forests in Poland. CWD diameter and number of sections i.e. number interconnected trunks and branches making up CWD were used as continuous variables, while taxa (deciduous vs coniferous), presence of additional decay class (present vs absent) and main decay class as categorical variables; decay class were used as dummy variables (1 if debris was in the main decay class or 0 if debris holds no the main decay class) with 1^st^ decay class set as a reference; study area (five levels) and plot (423 levels) identification numbers were used as random effects. The data was limited to debris identified as deciduous or coniferous taxa (N = 5809); see Methods for description of the study sites.

| Explanatory variable | Estimate | SE | z | Pr(>\|z\|) | SD |
| --- | --- | --- | --- | --- | --- |
| Intercept | -0.21 | 0.08 | -2.54 | 0.011 |  |
| CWD diameter | 0.03 | 0.00 | 49.46 | 0.000 |  |
| Number of sections | 0.08 | 0.01 | 11.68 | 0.000 |  |
| Taxa (deciduous) | 0.12 | 0.03 | 3.61 | 0.000 |  |
| 0<sup>th</sup> decay class (present) | 0.18 | 0.09 | 2.05 | 0.040 |  |
| 2<sup>nd</sup> decay class (present) | 0.21 | 0.03 | 6.11 | 0.000 |  |
| 3<sup>rd</sup> decay class (present) | 0.33 | 0.03 | 9.38 | 0.000 |  |
| 4<sup>th</sup> decay class (present) | 0.17 | 0.04 | 4.29 | 0.000 |  |
| 5<sup>th</sup> decay class (present) | -0.19 | 0.06 | -3.02 | 0.003 |  |
| Additional decay classes (present) | 0.28 | 0.04 | 7.05 | 0.000 |  |
| Random effect: Study area |  |  |  |  | 0.16 |
| Random effect: Plot identification |  |  |  |  | 0.22 |

## Discussion

To our knowledge, this is the first structured catalogue of DreMs that provides a systematic overview of microhabitats occurring on downed CWD. Such a standardized framework is crucial to make results comparable across studies and to enable DreMs to be considered alongside TreMs in biodiversity assessments (Kraus et al., 2016a; Larrieu et al., 2018). The richness and composition of DreMs recorded in our study are consistent with studies examining individual deadwood-associated structures in natural mixed and deciduous temperate forests of Central Europe. The frequency of bryophyte mats in our dataset (ca 60% of CWD) falls within the range of 48–97% reported by Ódor & van Hees (2004) in Hungarian beech forests. The frequency of polypore fungal fruiting bodies and annual fungi found in our study on coniferous taxa (21.6% and 11.3%, respectively) corresponds to frequency of macrofungal fruiting bodies found on large (100–150 cm in diameter) Norway spruce logs in primeval forest in Czechia (33%; Holec et. al., 2020). The frequency of cavities observed in our study (3.6%) corresponds within the value of 3.5% reported by Zumr et al. (2024) in beech forest in Czechia. However, the study of the entire assemblage of DreMs is not mirrored in available researches. Therefore, structured catalogue provided in present study can be used in future works on DreM richness and composition in temperate forests.

Our results indicate that size and decay class of CWD are linked with DreMs. We found that the DreM richness increased with CWD diameter, as larger debris provides more surface area (Kolényová et al., 2024), greater water-holding capacity (Přívětivý & Šamonil, 2021) and more heterogeneous microclimate (Phillips & Garrick, 2026), which together promotes both the colonization by fungi and bryophytes and the development of cavities and cracks (Renvall, 1995; Heilmann-Clausen & Christensen, 2003). Our results indicate that freshly fallen CWD (0^th^ decay class) supported relatively high DreM richness, and that DreM richness peaked at intermediate stages as decomposition progressed. The high DreM richness of freshly fallen CWD likely reflects the coexistence of structures inherited from living trees, such as lichen patches, vines or cracks, together with structures that develop immediately after tree fall, including uprooted root plates. Therefore, this specific and most commonly short term stage of transition between living standing tree and downed dead tree combines both TreMs and DreMs. Intermediate decay stages corresponds with earlier findings that wood in mid-decay stages is soft enough to allow the formation of new features, while still structurally stable enough to sustain a broad range of organisms (Ódor & van Hees, 2004; Kolényová et al., 2024; Holec et al., 2025). In contrast, advanced stages of decomposition were characterized by a decline in DreM richness, likely due to the disintegration of wood and disappearance of structures that develop earlier on less decomposed substrates. Moreover, both the complex architecture, i.e. number of trunk and branch sections composing a CWD, and the coexistence of multiple decay classes within single debris further enhanced DreM richness. Such mosaic-like structures create fine-scale heterogeneity that probably supports diverse groups of organisms and enable development of structures related to varying ecological conditions (Kolényová et al., 2024; Zumr et al., 2024). Thus, substrate heterogeneity arising from CWD size, decomposition and architecture emerges as a primary mechanism driving the diversity of DreMs.

DreM richness observed on deciduous CWD was higher than on coniferous. This pattern can be associated with more rapid decomposition of wood of deciduous trees, which can faster lead to higher substrate heterogeneity (Zhou et al., 2007). Deciduous trees typically exhibit greater trunk and branch architecture, including trunk forks and larger branching systems, which may promote, for example, the formation of cracks or cavities, dendrotelms or empty interiors (Ibach, 2005). Moreover, higher moisture recorded on deciduous taxa (Přívětivý & Šamonil, 2021) supports bryophytes, fungi and other moisture-dependent organisms (Andersson & Hytteborn, 1991; Ódor et al., 2006). Consequently, deciduous CWD provides more suitable conditions for formation of DreMs across a wide spectrum of taxa. In our study, silver birch or crack and white willows have high DreM richness, highlighting the importance of these early successional taxa for deadwood-associated fine-scale habitat heterogeneity (Przepióra & Ciach, 2023). Although coniferous CWD exhibited lower overall DreM richness, Norway spruce played an important role as substrate for specific DreM types, particularly woody plant cover, which, despite being a rare microhabitat overall (frequency < 5% of all CWD), occurred more frequent (3.9%) compared to other tree species (0.2–1.6%). These results emphasize the important role of tree species diversity for forest biodiversity and underscore the need for broad tree species representation in temperate forests (Ampoorter et al., 2019; Messier et al., 2022).

Although our study did not directly assess the functional role of DreMs for particular organism groups, numerous studies have demonstrated that structures developed on CWD, e.g. such as hollow logs, loose bark, fungal fruiting bodies, bryophyte mats or uprooted trees, are widely used by forest-dwelling organisms, including insects, amphibians, birds and mammals (Wesołowski, 1983; Bull & Heater, 2000; Bull, 2002; Fritts et al., 2015). This body of evidence suggests that DreMs may contribute substantially to the maintenance of forest biodiversity and ecosystem functioning, though, in-depth understanding of the mechanisms linking DreMs with specific organisms or species communities requires further research. As many saproxylic organisms have limited dispersion potential and the distribution of DreMs at the stand and landscape level may influence their persistence, investigation of spatial and temporal continuity of DreMs is of great importance. Given DreM frequency and diversity, these features have considerable potential to serve as indicators of forest naturalness, much like TreMs are increasingly used in habitat quality assessments.

## Conclusion

Our study shows that CWD in old-growth temperate forests hosts numerous and diversified assemblage of DreMs. The DreM richness was associated with characteristic of CWD, i.e. diameter, decomposition stage and debris architecture. In particular, large CWD, deciduous species, domination of intermediate decay classes and presence of additional decay classes co-occurring within single debris supported the highest DreM richness. By developing the comprehensive catalogue of DreMs and applying it in the field surveys conducted in the best-preserved old-growth forest in Poland, we demonstrated that, besides traditional descriptors such as volume or stage of decomposition, heterogeneous deadwood-associated features could be included in CWD quality assessment. By focusing on forests characterized by long ecological continuity and minimal human disturbance, our results also provide a reference for DreMs in temperate forests. Our work complements the existing concept of TreMs and expands current approaches to forest biodiversity assessment by incorporating fine-scale habitat structures associated specifically with decomposing downed woody material.

## Acknowledgements

This study was financially supported by the National Science Centre in Poland through the Preludium grant (2021/41/N/NZ9/03441), the Opus grant (2024/55/I/NZ8/02986) and by the Ministry of Science and Higher Education of the Republic of Poland within the framework of statutory funds awarded to the Faculty of Forestry, University of Agriculture in Kraków. FP was supported by the Foundation for Polish Science. We thank the management of Tatra National Park, Bieszczady National Park, Świętokrzyski National Park and Białowieża National Park, as well as the Regional Directorate for Environmental Protection in Warsaw, for granting access to the study areas and permission to conduct the study. We are grateful to Jakub Morawski and Paweł Lewandowski for their assistance during the fieldwork.

## Ethical statement

The study was performed in accordance with Polish law.

## Data availability

The data used in this study is available on request from the corresponding author.

## Supplementary materials

**Table S1.**
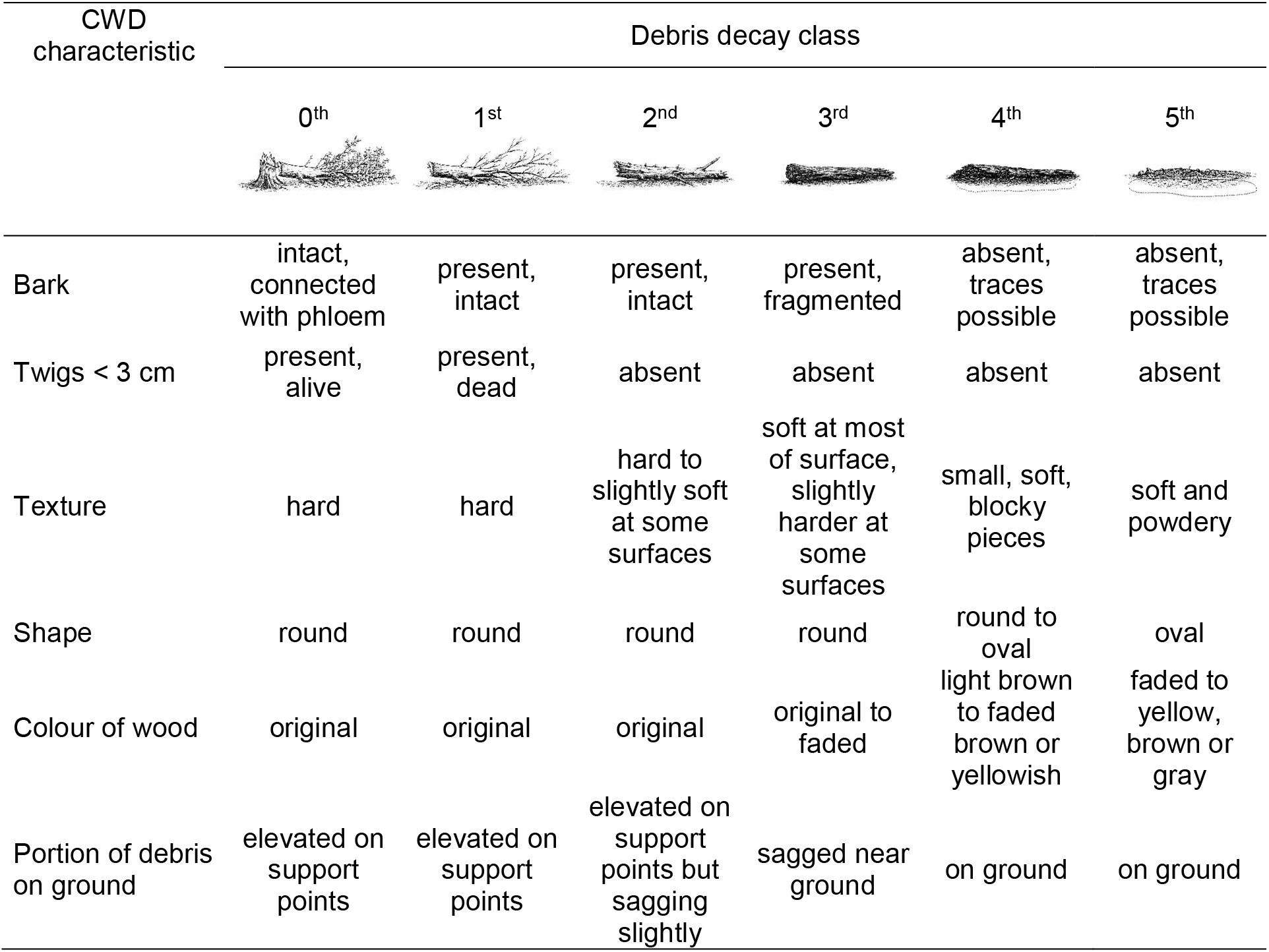
Classification of downed coarse woody debris (CWD) according to decay classes based on Maser et al. (1979), modified by the additional decay class (0^th^) representing recently fallen trees that remained connected to living tissues through intact phloem.

| CWD characteristic | Debris decay class |  |  |  |  |  |
| --- | --- | --- | --- | --- | --- | --- |
|  | 0<sup>th</sup> | 1<sup>st</sup> | 2<sup>nd</sup> | 3<sup>rd</sup> | 4<sup>th</sup> | 5<sup>th</sup> |
| Bark | intact, connected with phloem | present, intact | present, intact | present, fragmented | absent, traces possible | absent, traces possible |
| Twigs < 3 cm | present, alive | present, dead | absent | absent | absent | absent |
| Texture | hard | hard | hard to slightly soft at some surfaces | soft at most of surface, slightly harder at some surfaces | small, soft, blocky pieces | soft and powdery |
| Shape | round | round | round | round | round to oval | oval |
| Colour of wood | original | original | original | original to faded | light brown to faded brown or yellowish | faded to yellow, brown or gray |
| Portion of debris on ground | elevated on support points | elevated on support points | elevated on support points but sagging slightly | sagged near ground | on ground | on ground |

**Table S2.** Number of downed coarse woody debris of **(a)** deciduous, **(b)** coniferous and **(c)** unknown taxa, found in old-growth forests in Poland. Study sites included Norway spruce *Picea abies*-dominated upper montane zone forests protected as Tatra National Park (TNP), European beech *Fagus sylvatica*-dominated lower montane zone forests protected as Bieszczady National Park (BdNP), willow *Salix* sp.-poplar *Populus* sp. riparian forest of Middle Vistula Valley (MVV), mixed European beech-silver fir *Abies alba* mountain forest protected as Świętokrzyski National Park (ŚNP) and pedunculate oak *Quercus robur*-small-leaved lime *Tilia cordata*-European hornbeam *Carpinus betulus* forest protected as Białowieża National Park (BwNP).

| Taxa |  | All areas pooled | TNP | BdNP | MVV | ŚNP | BwNP |
| --- | --- | --- | --- | --- | --- | --- | --- |
| **(a) Deciduous** |  |  |  |  |  |  |  |
| European beech | *Fagus sylvatica* | 1474 |  | 1439 |  | 35 |  |
| Unknown deciduous species |  | 854 |  |  | 457 | 34 | 363 |
| Crack willow and white willow | *Salix fragilis* and *Salix alba* | 400 |  |  | 400 |  |  |
| European hornbeam | *Carpinus betulus* | 311 |  |  |  |  | 311 |
| White poplar and black poplar | *Populus alba* and *Populus nigra* | 166 |  |  | 166 |  |  |
| Small-leaved lime | *Tilia cordata* | 79 |  |  |  |  | 79 |
| Rowan | *Sorbus aucuparia* | 79 | 2 |  |  | 77 |  |
| Pedunculate oak | *Quercus robur* | 75 |  |  |  |  | 75 |
| European aspen | *Populus tremula* | 70 |  | 27 | 1 | 15 | 27 |
| European ash | *Fraxinus excelsior* | 46 |  |  |  |  | 46 |
| Silver birch | *Betula pendula* | 30 |  | 5 |  | 21 | 4 |
| Goat willow | *Salix caprea* | 73 | 5 | 68 |  |  |  |
| Common hazel | *Corylus avellana* | 19 |  |  |  |  | 19 |
| Norway maple | *Acer platanoides* | 14 |  |  |  |  | 14 |
| Boxelder maple | *Acer negundo* | 8 |  |  | 8 |  |  |
| Elms | *Ulmus* sp. | 2 |  |  | 1 |  | 1 |
| European pear | *Pyrus communis* | 1 |  |  | 1 |  |  |
| **(b) Coniferous** |  |  |  |  |  |  |  |
| Norway spruce | *Picea abies* | 1984 | 1617 | 3 |  | 20 | 344 |
| Silver fir | *Abies alba* | 122 |  |  |  | 122 |  |
| Scotch pine | *Pinus sylvestris* | 2 |  |  |  | 2 |  |
| **(c) Unknown** |  |  |  |  |  |  |  |
| Unknown taxa |  | 194 | 0 | 0 | 34 | 34 | 126 |

**Table S3.** Frequency (percentage) of downed coarse woody debris (CWD) in specific decay class found in old-growth forests in Poland, calculated for taxa with N ≥ 30 (see Table S2 for number of surveyed CWD): all CWD pooled, deciduous taxa pooled, coniferous taxa pooled, Norway spruce *Picea abies*, European beech *Fagus sylvatica*, crack willow *Salix fragilis* and white willow *Salix alba* pooled, European hornbeam *Carpinus betulus*, white poplar *Populus alba* and black poplar *Populus nigra* pooled, silver fir *Abies alba*, small-leaved lime *Tilia cordata*, rowan *Sorbus aucuparia*, pedunculate oak *Quercus robur*, goat willow *Salix caprea*, European aspen *Populus tremula*, European ash *Fraxinus excelsior* and silver birch *Betula pendula*.

| Taxa | Decay class |  |  |  |  |  |
| --- | --- | --- | --- | --- | --- | --- |
|  | 0<sup>th</sup> | 1<sup>st</sup> | 2<sup>nd</sup> | 3<sup>rd</sup> | 4<sup>th</sup> | 5<sup>th</sup> |
| Total | 1.0 | 7.7 | 46.0 | 30.2 | 11.7 | 3.4 |
| Deciduous | 1.4 | 8.9 | 41.4 | 34.8 | 10.9 | 2.6 |
| Coniferous | 0.2 | 6.1 | 57.3 | 22.4 | 11.2 | 2.8 |
| Norway spruce | 0.1 | 6.1 | 59.4 | 21.4 | 10.4 | 2.6 |
| European beech | 0.6 | 9.8 | 40.6 | 31.2 | 13.5 | 4.2 |
| Crack willow and white willow | 6.2 | 17.8 | 47.5 | 19.8 | 8.2 | 0.5 |
| European hornbeam | 0.6 | 8.7 | 39.9 | 36.0 | 13.5 | 1.3 |
| White poplar and black poplar | 1.8 | 13.9 | 53.0 | 22.3 | 6.0 | 3.0 |
| Silver fir | 2.5 | 5.7 | 23.0 | 37.7 | 24.6 | 6.6 |
| Small-leaved lime | 7.6 | 11.4 | 43.0 | 30.4 | 7.6 | 0.0 |
| Rowan | 2.5 | 3.8 | 54.4 | 35.4 | 2.5 | 1.3 |
| Pedunculate oak | 0.0 | 2.7 | 52.0 | 45.3 | 0.0 | 0.0 |
| Goat willow | 0.0 | 1.4 | 16.4 | 78.1 | 4.1 | 0.0 |
| European aspen | 0.0 | 5.7 | 40.0 | 47.1 | 7.1 | 0.0 |
| European ash | 2.2 | 28.3 | 58.7 | 10.9 | 0.0 | 0.0 |
| Silver birch | 0.0 | 0.0 | 26.7 | 66.7 | 6.7 | 0.0 |

**Table S4.** Frequency (percentage) of downed coarse woody debris (CWD) in a particular decay class that holds additional fragments in other decay class found in old-growth forests in Poland.

| Description | Frequency |  |  |
| --- | --- | --- | --- |
|  | All CWD<br>(N = 6003) | Deciduous taxa<br>(N = 3702) | Coniferous taxa<br>(N = 2017) |
| 0<sup>th</sup> downed living tree | 0.0 | 0.0 | 0.0 |
| 1<sup>st</sup> decay class | 0.0 | 0.0 | 0.0 |
| 2<sup>nd</sup> decay class | 0.2 | 0.1 | 0.4 |
| 3<sup>rd</sup> decay class | 1.5 | 0.2 | 3.8 |
| 4<sup>th</sup> decay class | 1.0 | 0.4 | 2.0 |
| 5<sup>th</sup> decay class | 0.3 | 0.1 | 0.6 |

## Appendix. 1.

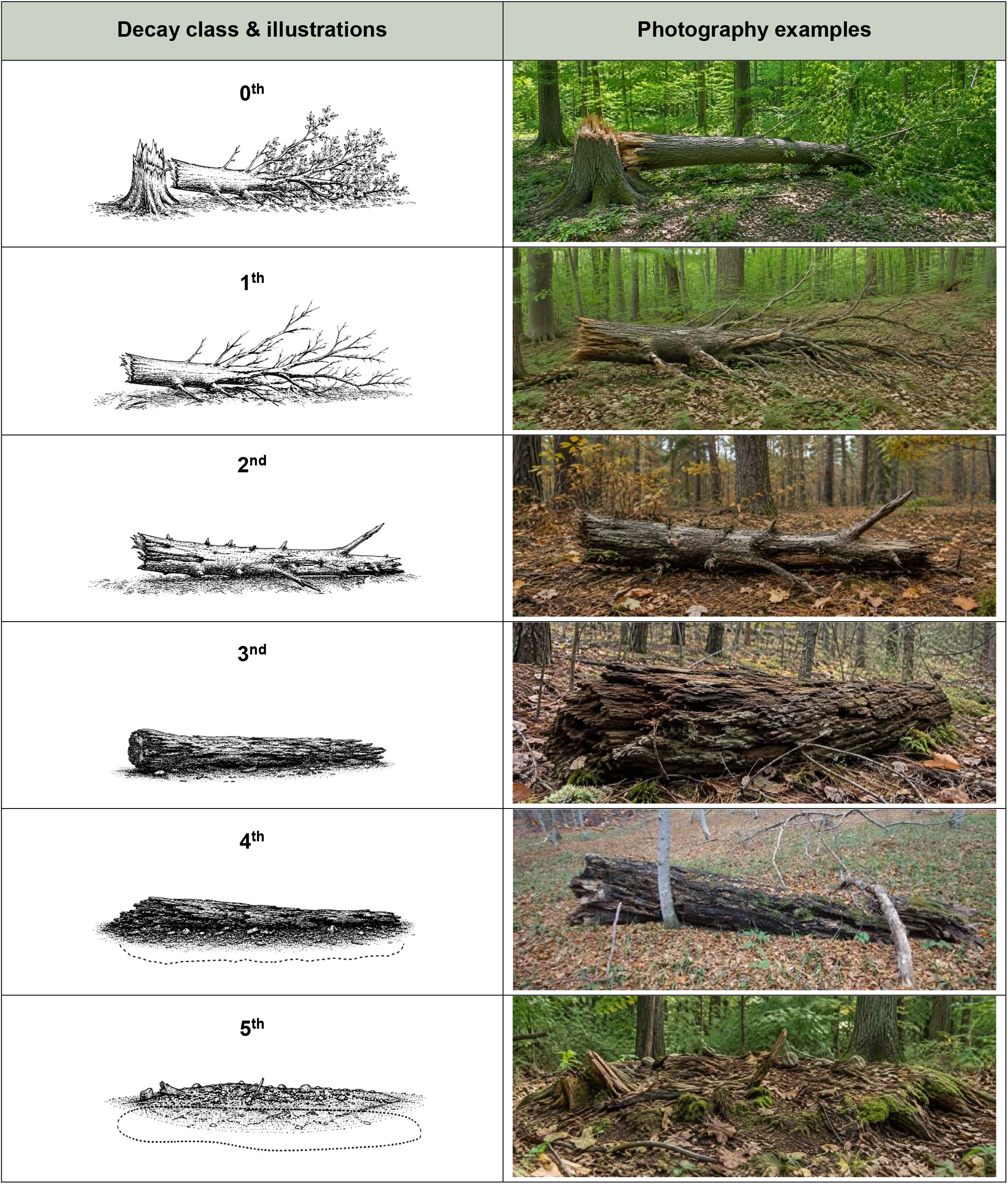

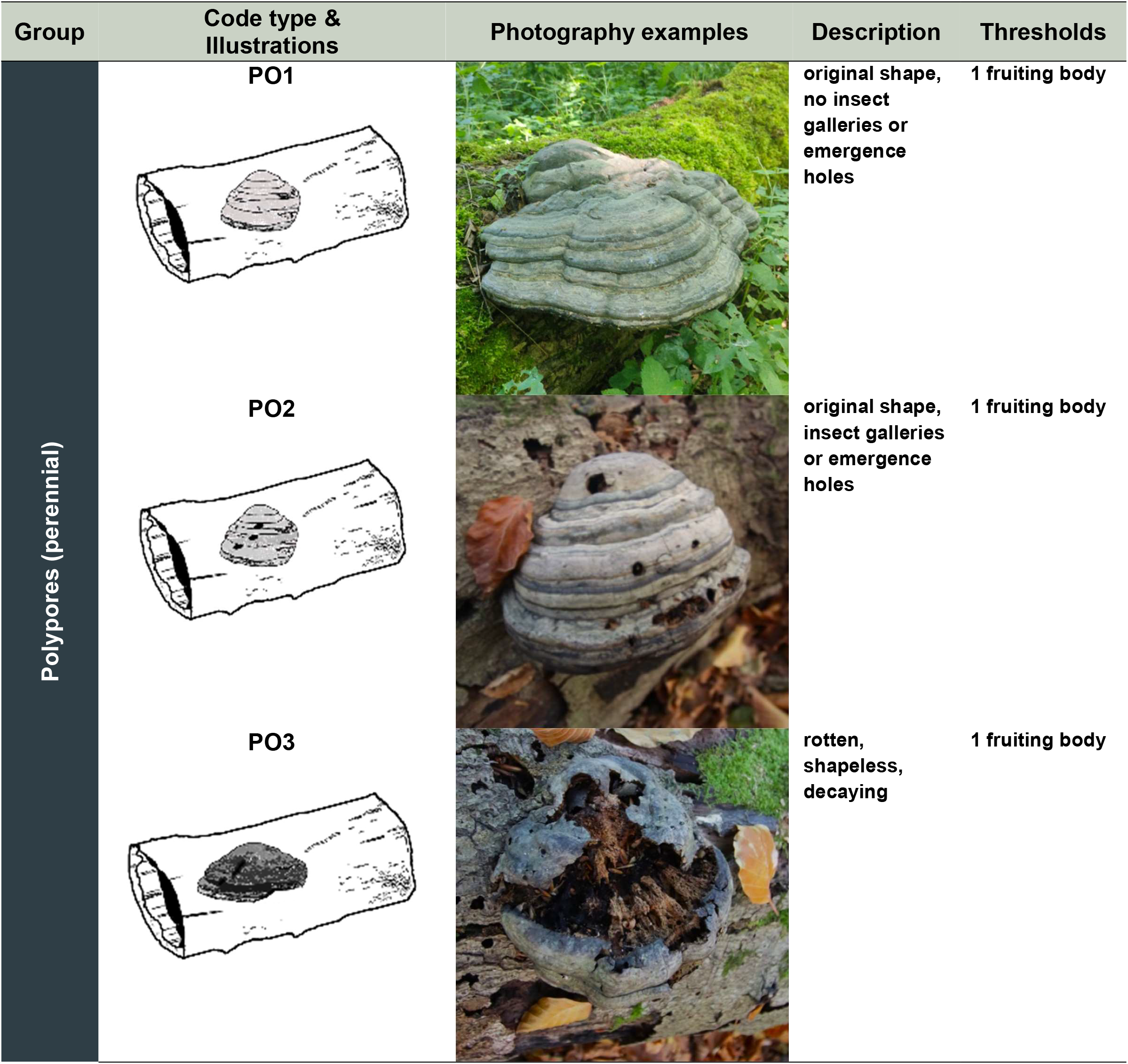

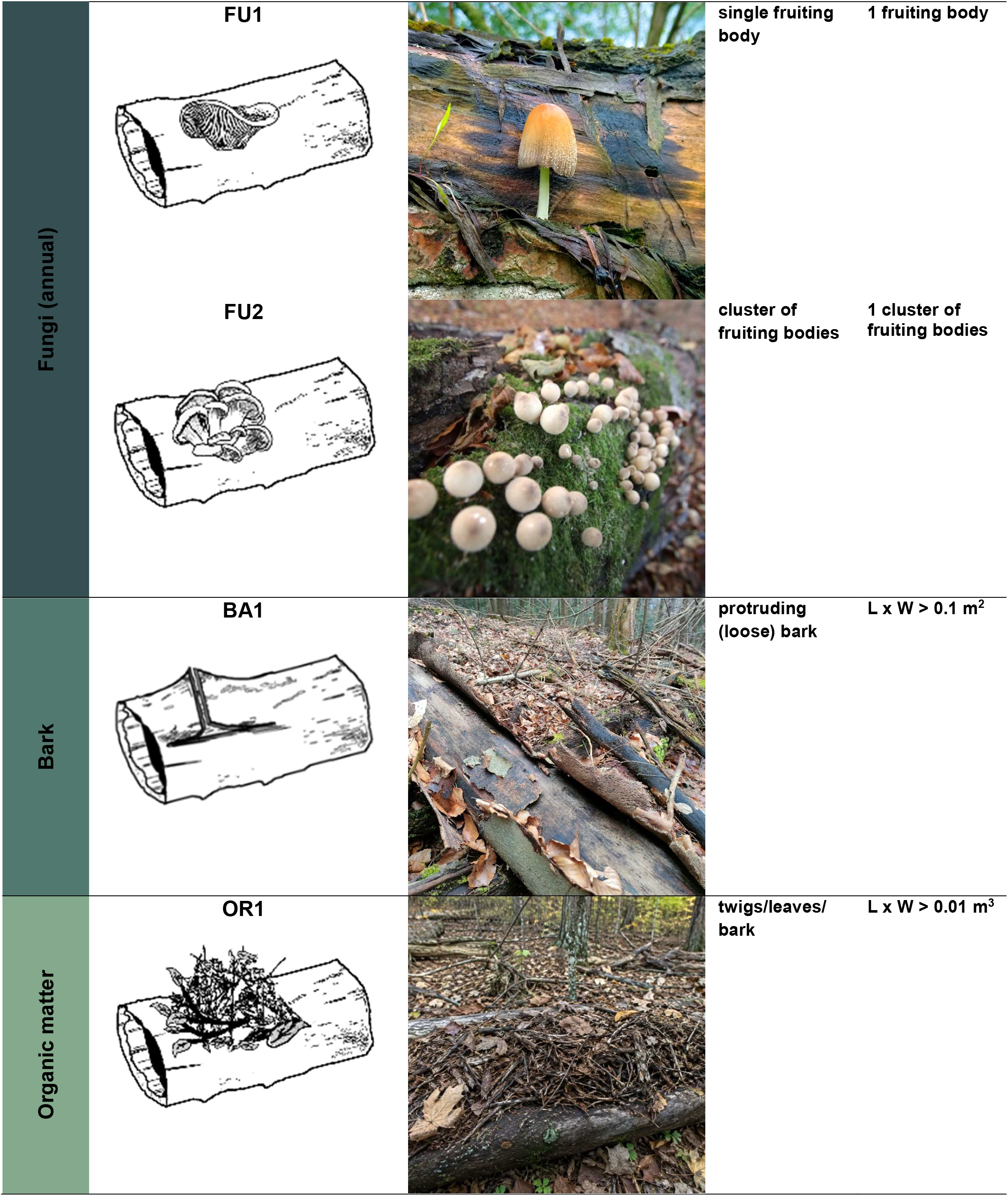

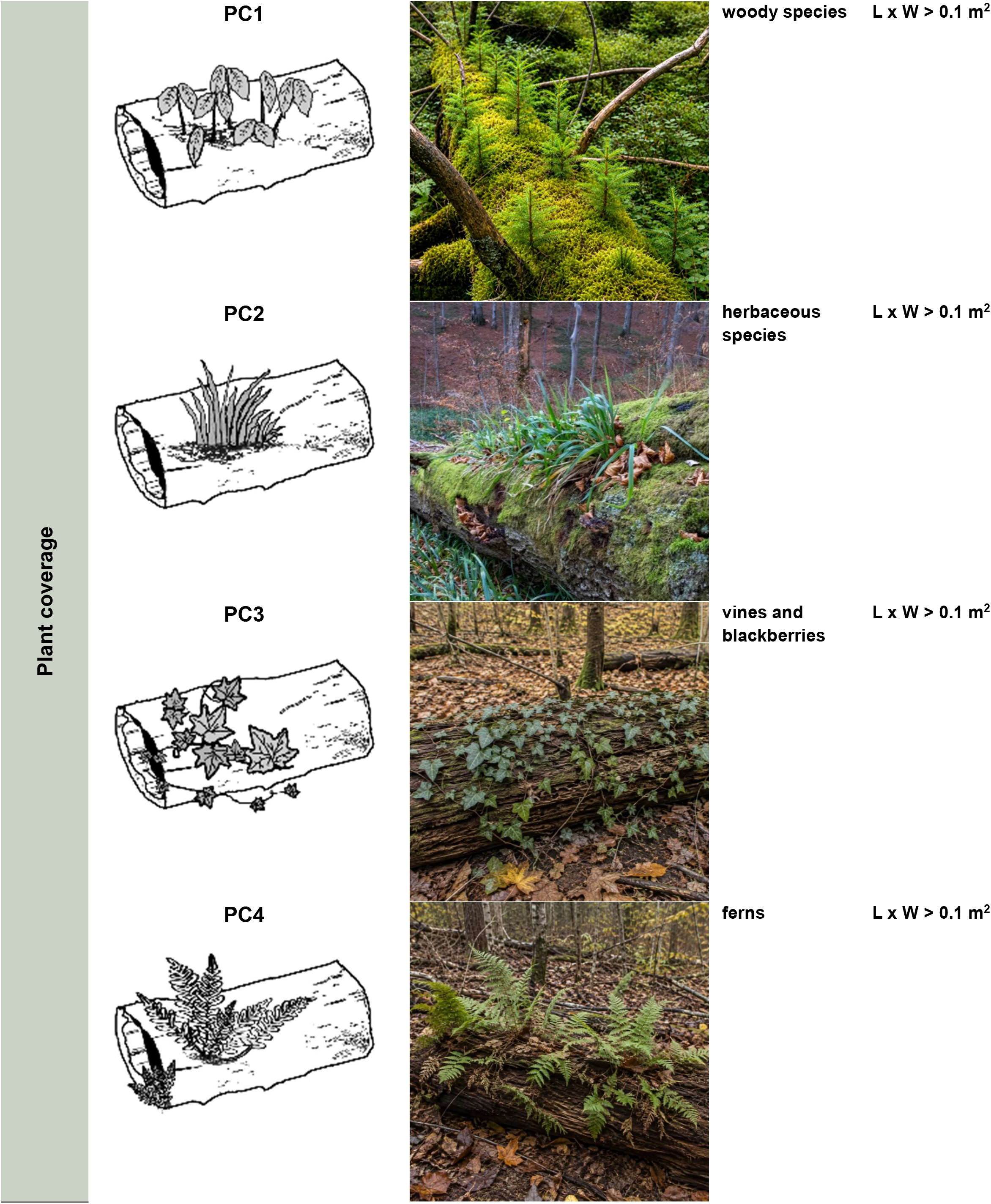

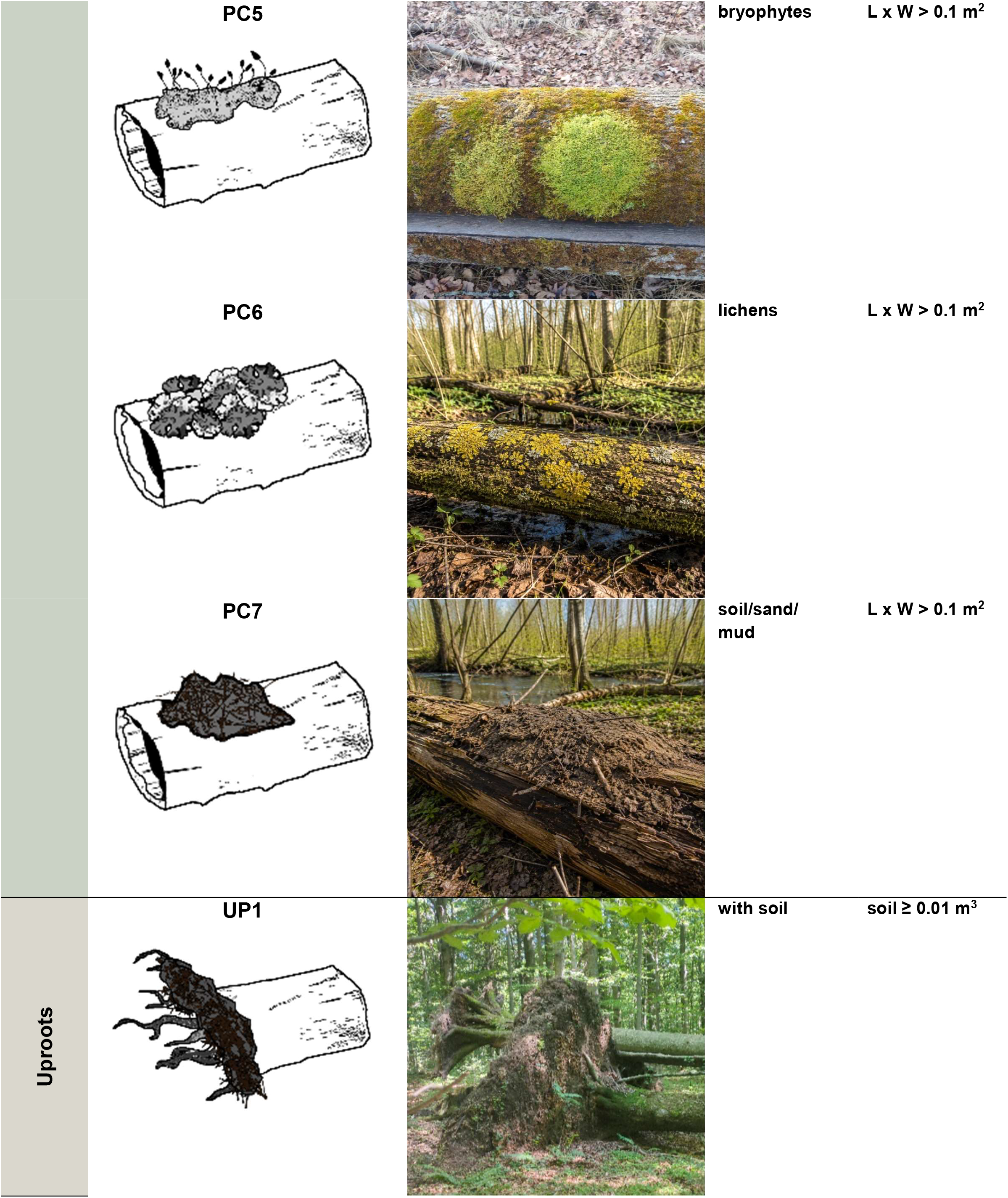

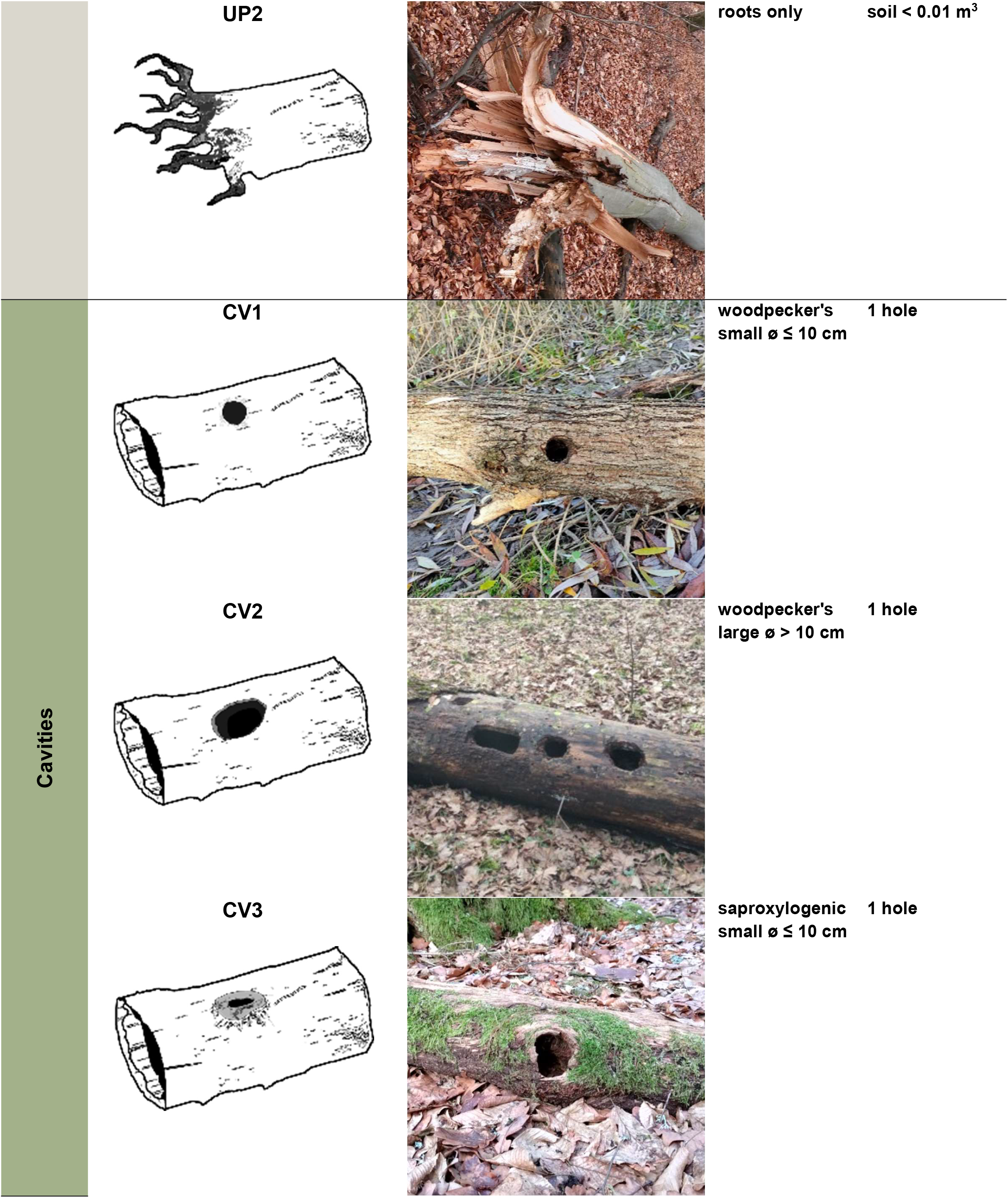

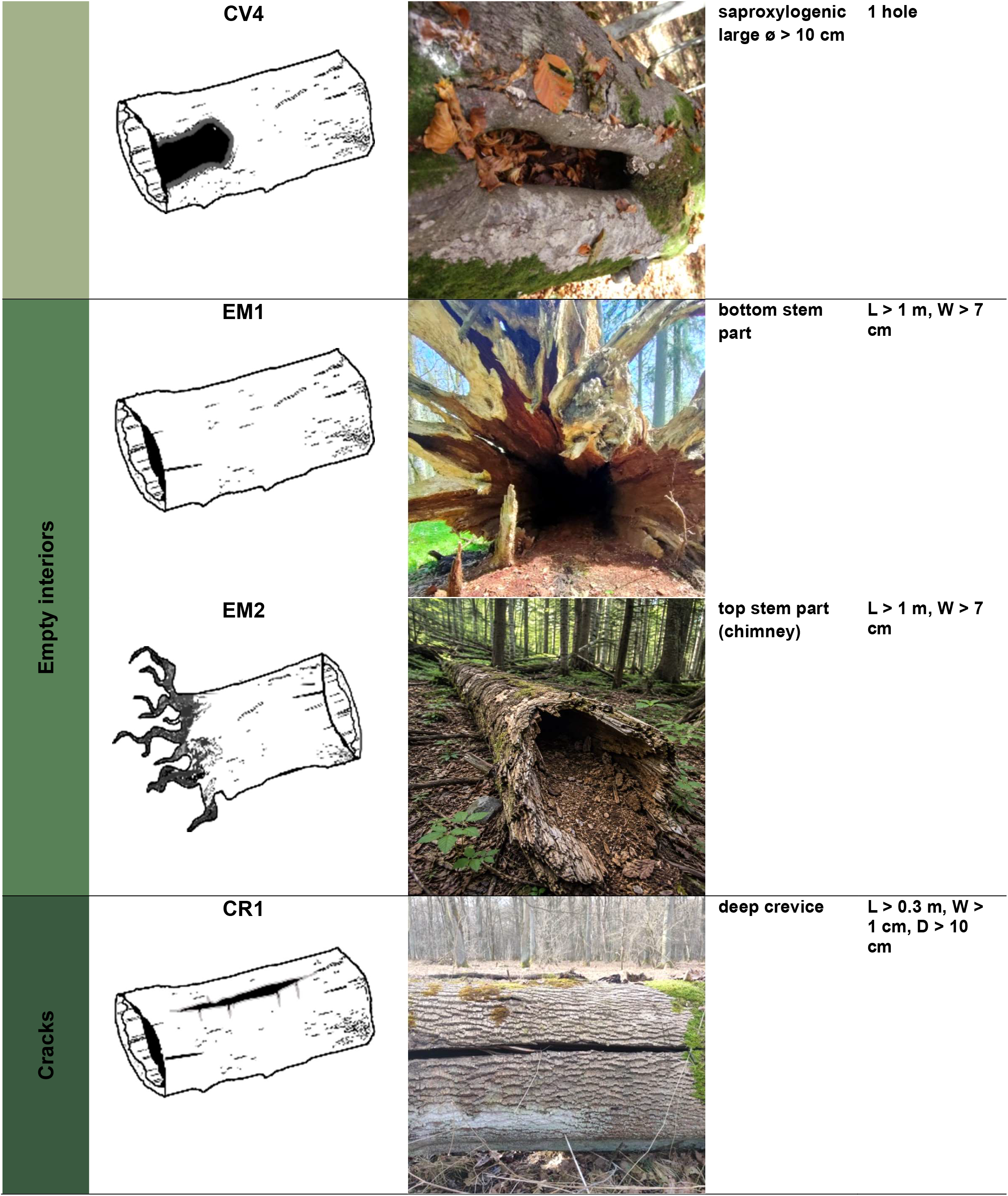

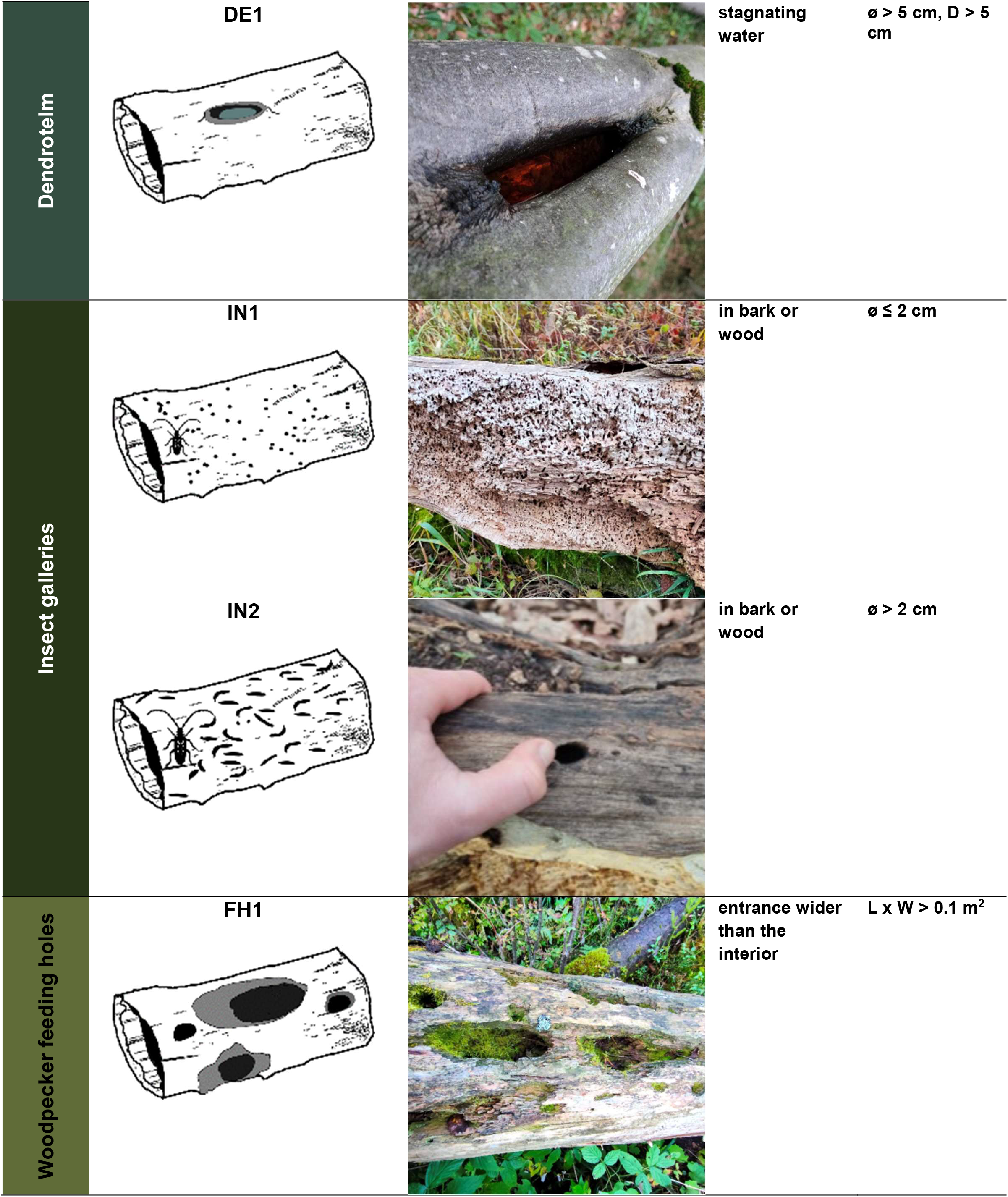

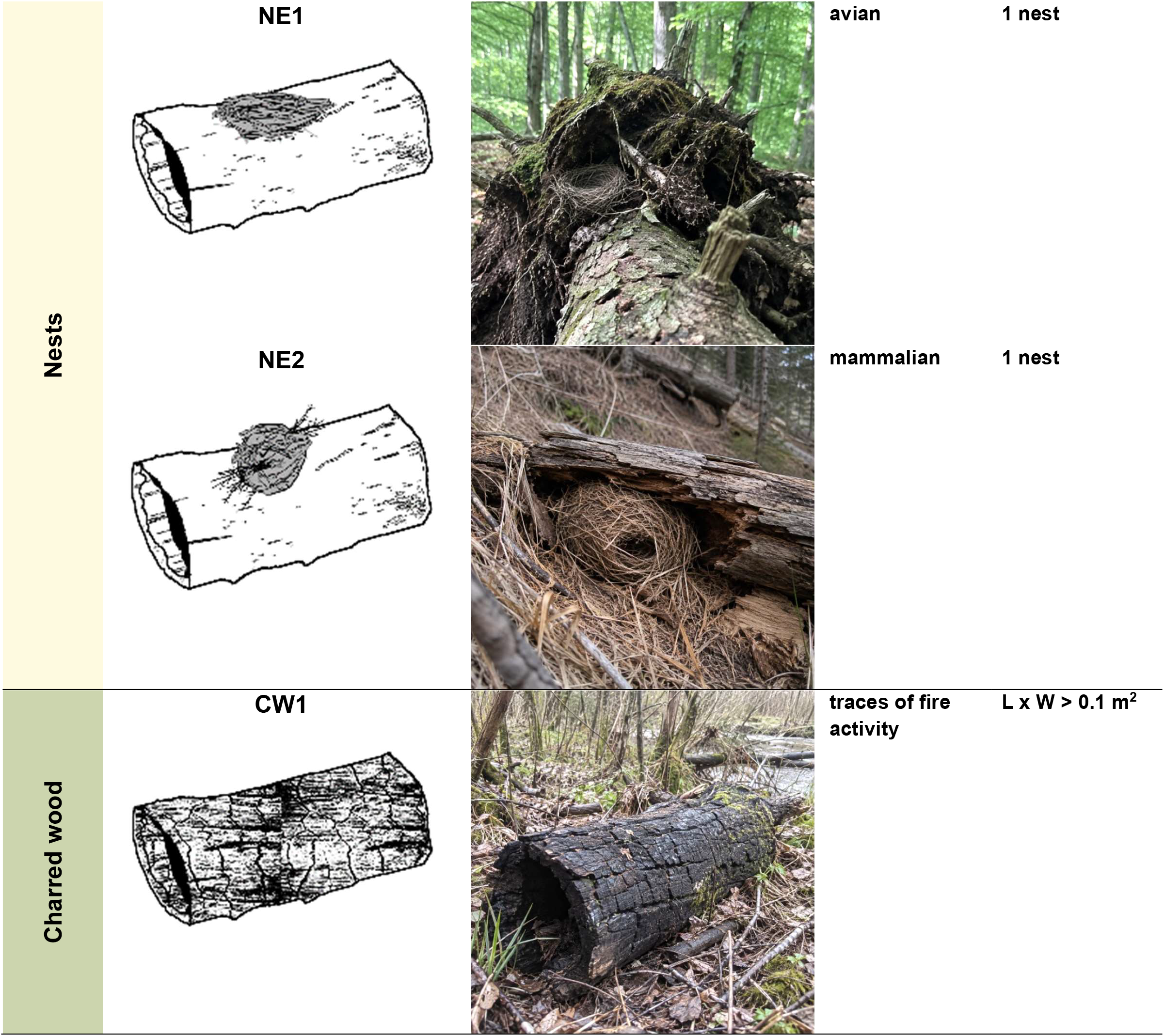

